# Mast cells participate in the development of diastolic dysfunction in diabetic obese mice

**DOI:** 10.1101/2020.06.16.154872

**Authors:** Sarah Guimbal, Lauriane Cornuault, Pierre-Louis Hollier, Candice Chapouly, Marie-Lise Bats, Alain-Pierre Gadeau, Thierry Couffinhal, Marie-Ange Renault

## Abstract

**Rational:** Heart failure with preserved ejection fraction (HFpEF) is a growing epidemiologic issue. However, to date, its pathophysiology remains poorly understood.

**Objective:** Our goal was to investigate the role of microvessel disease in the pathophysiology of diastolic dysfunction.

**Findings:** To do so, we used Leptin receptor deficient (Lepr^db/db^) female mice as a model of diastolic dysfunction. In these mice, the increased end diastolic pressure (EDP) signing diastolic dysfunction is associated with vascular leakage, endothelial cell activation and leucocyte infiltration. Strikingly, a RNA sequencing analysis of the cardiac vascular fraction of both Lepr^db/db^ and control female mice confirmed endothelial dysfunction and systemic inflammation but also revealed a strong increase in several mast cell markers (notably FceR1a, Tryptase and Chymase). We then histologically confirmed an accumulation of activated mast cells in the heart of Lepr^db/db^ mice. Importantly, mast cell degranulation inhibition reduced EDP, vascular leakage and leucocyte infiltration in Lepr^db/db^ mice.

**Conclusion:** Mast cells play a critical role in the development of cardiac microvessel disease and diastolic dysfunction.

## Introduction

A significant proportion of patients with clinical syndrome of heart failure (HF) happens to have a normal ventricular ejection fraction (EF) referred to as heart failure with preserved ejection fraction (HFpEF) as opposed to patients with reduced ejection fraction (HFrEF) ^1^. HFpEF currently accounts for about 50% of all HF patients, and its prevalence relative to HFrEF is rising at a rate of approximately 1% per year. HFpEF patients are generally older, more often female and have a high prevalence of cardiovascular and non-cardiovascular comorbidities, such as obesity, metabolic syndrome, diabetes mellitus type 2, hypertension, atrial fibrillation or renal dysfunction ^1^. HFpEF is diagnosed in the presence of HF signs and/or symptoms, preserved systolic left ventricular (LV) function, with an LV ejection fraction >50% and LV end-diastolic volume index <97 ml/m^2^ with evidence of diastolic LV dysfunction ^2^. In recent years, there have been significant advances in understanding aetiologies and mechanisms of HFrEF, leading to new therapeutic options. In contrast, if some progresses on HFpEF pathophysiology have been made, therapeutic targets are still missing and no therapies are available to reduce hospitalization or mortality for HFpEF patients. There is, therefore, an urgent need for a better understanding of HFpEF pathophysiology in order to develop new drugs ^3^. A unifying, however untested, theory of the pathophysiology of HFpEF suggests that comorbidities lead to a systemic low grade inflammation, which triggers coronary microvascular dysfunction ^2,4–6^ and subsequently heart failure.

Endothelial dysfunction mostly refers to impaired endothelium-dependent, nitric oxide-mediated vasorelaxation. Nevertheless, endothelial dysfunction actually regroups an ensemble of features that include acquirement of a pro-inflammatory /pro-thrombotic phenotype due to an increased expression of adhesion and pro-thrombotic molecules such as vascular cell adhesion molecule-1 (VCAM-1) and intercellular adhesion molecule-1 (ICAM-1), abnormal vascular leakage due to altered endothelial intercellular junctions and endothelial dedifferentiation promoting endothelial to mesenchymal transition ^7^. EC dysfunction may then lead to a compromised heart perfusion because of impaired NO-dependent vasodilatation, cardiac oedema or thrombi formation. Besides, it may also promote cardiac fibrosis and inflammation ^4,5,8^. Finally, an impaired production of endothelial-derived active substances as NO, endothelin-1, angiotensin II (AII), Neuregulin is also proposed to directly affect cardiomyocyte passive tension^9–11^.

What is known so far is that HFpEF is associated with an increased endothelial expression of ICAM-1 and E-selectin, an increased oxidative stress and a decreased eNOS activity both in animal models and in patients ^12^. Also, brachial artery flow-mediated dilation assessment and laser Doppler flow measurement have revealed an impaired vascular and microvascular function in patients with HfpEF 5. Finally, HFpEF is typically associated with increased myocardial stiffening (including increased cardiomyocyte passive stiffness and accumulation of stiff and resistant collagen fibres). Nevertheless the causal role of endothelial dysfunction in the pathophysiology of HFpEF has never been established.

In the present study we used Lepr^db/db^ female mice as a model of HFpEF to further characterize the phenotype and mechanisms of cardiac microvessel dysfunction associated with diastolic dysfunction. Notably, Lepr^db/db^ female mice recapitulate the main risk factors for HFpEF i.e. obesity, diabetes and female gender. As a major breakthrough, this study demonstrates for the first time that abnormally increased mast cell activation participates in the pathophysiology of both cardiac microvessel disease and diastolic dysfunction in Lepr^db/db^ female mice.

## Methods

### Mice

Lepr^db^ mice (BKS.Cg-*Dock7*^*m*^/+ *Lepr*^*db*^/+J) were obtained from Charles River laboratories and bred together to obtain Lepr^db/db^ and control Lepr^db^/+ mice.

Animal experiments were performed in accordance with the guidelines from Directive 2010/63/EU of the European Parliament on the protection of animals used for scientific purposes and approved by the local Animal Care and Use Committee of Bordeaux University. Only females were used.

### Cromolyn Sodium therapy

To prevent mast cell degranulation, mice were treated with 50 mg/kg/day cromolyn sodium (Abcam) via intra-peritoneal injections for 28 days. Untreated mice received 0,9% NaCl daily intra-peritoneal injections.

### Blood tests/NFS

Blood samples were collected on heparin retroorbital bleeding. Blood cell counts were determined using an automated counter (scil Vet abc Plus+). Plasmas were separated by a 10 min centrifugation at 2500 g and stored at -80°C. Concentrations of the following biochemical markers were measured using an Architect CI8200 analyzer (Abbott Diagnostics, North Chicago, Illinois, USA): triglycerides by the lipoprotein-lipase/glycerol kinase/oxidase/peroxidase method; total cholesterol by the esterase/oxidase/peroxidase method and HDL cholesterol by the accelerator/selective detergent/esterase/oxidase/peroxidase method. LDL cholesterol was then estimated using the Friedewald formula (LDL cholesterol (mmol/L) = total cholesterol – HDL cholesterol – (triglycerides/2,2) or LDL cholesterol (mg/dL) = total cholesterol – HDL cholesterol – (triglycerides/5)).

### Echocardiography

Left-ventricular ejection fraction and LV dimension will be measured on a high-resolution echocardiographic system equipped with a 30-MHz mechanical transducer (VEVO 2100, VisualSonics Inc.) as previously described ^31,32^. Mice were anchored to a warming platform in a supine position, limbs were taped to the echocardiograph electrodes, and chests were shaved and cleaned with a chemical hair remover to minimize ultrasound attenuation. UNI’GEL ECG (Asept Inmed), from which all air bubbles had been expelled, was applied to the thorax to optimize the visibility of the cardiac chambers. Ejection fractions were evaluated by planimetry as recommended (Schiller et al. 1989). Two-dimensional, parasternal long-axis and short-axis views were acquired, and the endocardial area of each frame was calculated by tracing the endocardial limits in the long-axis view, then the minimal and maximal areas were used to determine the left-ventricular end-systolic (ESV) and end-diastolic (EDV) volumes, respectively. The system software uses a formula based on a cylindrical-hemiellipsoid model (volume=8.area^2^/3π/ length) ^33^. The left-ventricular ejection fraction was derived from the following formula: (EDV-ESV)/EDV*100. The cardiac wall thickness (Left ventricular posterior wall (LVPW), Inter-ventricular septum (IVS) and left ventricular internal diameter (LVID) were calculated by tracing wall limits in both the long and short axis views.

### LV pressure /systolic blood pressure measurement

LV diastolic pressure measurement was assessed using pressure–volume conductance catheter technique. Briefly, mice will be anesthetized with Isoflurane. A Scisense pressure catheter (Transonic) will be inserted into the LV through the common carotid artery. Pressure will be recorded using LabChart software. End diastolic pressure, dP/dt minimum and maximum, Tau and heart rate were automatically calculated by a curve fit through end-systolic and end-diastolic points on the pressure plot.

### Immuno-histological assessments

Prior to staining, heart were stopped in diastole using KCl, perfused and then fixed in 10% formalin for 4 hours, paraffin embedded and cut into 7 µm thick sections. Alternatively, heart were fresh frozen in OCT, then cut into 7 µm thick sections.

ECs were identified using rat anti-CD31 antibodies (Histonova, cat# DIA-310). Albumin was stained using sheep anti-albumin antibodies (Abcam, Cat# ab8940). Pan-leucocytes were identified using rat anti-mouse CD45 antibodies (BD Pharmingen Inc, Cat# 550539). Macrophages were identified using rat anti-CD68 antibodies (Biolegend, Cat# 137001). Neutrophils were identified using rat anti-Ly6G antibodies (BD Pharmingen Inc, Cat# 551459). T lymphocytes were identified using goat anti-CD3 antibodies (Santa Cruz Biotechnology, Inc, Cat# sc-1127). Mast cells were identified using goat anti-CD117 antibodies (R&D systems, Cat# AF1356). B-cells were stained using rat anti-B220/CD45R antibodies (R&D systems, Cat# MAB1217)

Fcer1a was identified using Armenian hamster anti-Fcer1a antibodies (eBioscience™, Cat# 14-5898-82). Cardiomyocytes were immunostained using rabbit anti-desmin antibodies (DB 148, Cat#DB148), mouse anti-sarcomeric actin antibodies (Sigma, Cat# A2172) and rabbit anti-Cx43 antibodies (Sigma, Cat# C6219)

For immunofluorescence analyzes, primary antibodies were resolved with Alexa Fluor^®^–conjugated secondary polyclonal antibodies (Invitrogen, Cat# A-21206, A-21208, A-11077, A-11057, A-31573, A-10037) and nuclei were counterstained with DAPI (1/5000). Negative controls using secondary antibodies only were done to check for antibody specificity.

Cardiomyocyte membranes were stained using Wheat Germ Agglutinin, Alexa Fluor™ 488 Conjugate (Invitrogen).

Fibrosis was quantified after Sirius red staining of heart sections by quantifying the percentage of red-positive area. Cardiomyocyte mean surface area was measured using the Image J software after membrane staining with Wheat Germ Agglutinin (WGA), Alexa Fluor™ 488 Conjugate (Invitrogen) in 5 pictures randomly taken under x630 magnification. Pictures were taken in areas where cardiomyocytes were oriented transversally. Capillary density (CD31+ vessels) was quantified in 4 pictures taken under x260 magnification in areas where cardiomyocytes were oriented transversally. To assess the mean capillary diameter, the diameter of 10 capillary randomly chosen in each picture was measured via Image J. Inflammatory cell density (CD45+, CD68+, CD3+ and GR1+) was quantified in 8 pictures randomly taken under x260 magnification. Mast cell density and mast cell degranulation was assessed after Toluidine blue staining of heart sections. Mast cells were counted in the entire section.

All pictures and quantifications (done using ImageJ/Fiji v2.0.0-rc-59 software (National Institute of Health, USA)) were performed by a blinded investigator. More precisely, all samples were assigned a random number prior to animal sacrifice, data collection and analysis. At the end of the experiment, the genotype/treatment for each animal was unveiled to allow data comparison and experimental conclusion.

### Isolation of mouse cardiac vascular fraction

Heart were dissociated using 2 mg/mL type IV collagenase (Gibco™, ThermoFisher) for 1 hour at 37°C and the resulting dissociated cells were filtrated on a 30 µm strainer. ECs cells were labelled with rat anti-mouse CD31 microbeads (Miltenyi Biotec). Labelled ECs and attached cells were then isolated magnetically. Importantly, collagenase digestion is incomplete and the CD31+ positive fraction included about 70% CD31+ endothelial cells, 29% NG2+ pericytes and 1% CD45+ leucocytes. The CD31+ fraction was referred as cardiac vascular fraction in the entire manuscript.

### Quantitative RT-PCR

RNA was isolated using Tri Reagent^®^ (Molecular Research Center Inc) as instructed by the manufacturer, from heart tissue that had been snap-frozen in liquid nitrogen and homogenized. For quantitative RT-PCR analyses, total RNA was reverse transcribed with M-MLV reverse transcriptase (Promega) and amplification was performed on an AriaMx Real Time PCR system (Agilent Technologies) using B-R SYBER^®^ Green SuperMix (Quanta Biosciences). Primer sequences are reported in Supplementary table 1.

The relative expression of each mRNA was calculated by the comparative threshold cycle method and normalized to 18S rRNA expression.

### Soluble/insoluble collagen measurement

Total cardiac collagen and insoluble cardiac collagen were quantified using the Sircol™ Soluble Collagen Assay and The Sircol™ INSOLUBLE Collagen Assay (Biocolor) according to the manufacturer’s instructions.

### GMPc quantification

GMPc was quantified using the Abnova™ Cyclic GMP Complete ELISA Kit (Thermo Fischer Scientific) according to the manufacturer’s instructions.

### RNA sequencing

RNA was isolated using Tri Reagent^®^ (Molecular Research Center Inc) as instructed by the manufacturer from the cardiac vascular fraction of Lepr^db/db^ mice and control Lepr^db/+^ mice. mRNA library preparation were realized following manufacturer’s recommendations (KAPA mRNA HyperPrep Kit from ROCHE). Final samples pooled library prep were sequenced on Nextseq 500 ILLUMINA, corresponding to 2×30Millions of reads per sample after demultiplexing.

Quality of raw data has been evaluated with FastQC ^34^. Poor quality sequences has been trimmed or removed with Trimmomatic ^35^ software to retain only good quality paired reads. Star v2.5.3a ^36^has been used to align reads on mm10 reference genome using standard options. Quantification of gene and isoform abundances has been done with rsem 1.2.28, prior to normalisation with edgeR bioconductor package ^37^. Finally, differential analysis has been conducted with the glm framework likelihood ratio test from edgeR. Multiple hypothesis adjusted p-values were calculated with the Benjamini-Hochberg procedure to control FDR. The FPKM values of all transcripts of which expression was significantly different between both groups in Supplemental table 2.

### Western blot analysis

Plamatic IgE level was evaluated by SDS PAGE using rat anti-mouse IgE antibodies (R&D systems Cat# MAB9935).

### Statistics

Results are reported as mean ± SEM. Comparisons between groups were analyzed for significance with the non-parametric Mann-Whitney test or a 2 way ANOVA followed by Sidak’s multiple comparison test (for kinetics analyses) using GraphPad Prism v8.0.2 (GraphPad Inc, San Diego, Calif). Differences between groups were considered significant when p≤0.05 (*: p≤0.05; **: p≤0.01; ***: p≤0.001).

## Results

### Lepr^db/db^ female mice have diastolic dysfunction from 3 months of age

First we quantified and characterized the kinetics of appearance of cardiovascular risk factors in the Lepr^db/db^ female mice and in their control Lepr^db^/+ littermates bred in our laboratory. Lepr^db/db^ female mice are significantly overweight from 6 weeks of age while diabetes appears from 3 months of age. Plasmatic HDL cholesterol is elevated from 6 weeks of age while LDL cholesterol is only increased after 6 months of age. Plasmatic triglycerides are elevated from 3 month of age. Besides Lepr^db/db^ female mice have mild hypertension from 3 month of age and display significantly increased circulating leucocytes (monocytes and neutrophils) later in life at 1 year of age (Supplemental Figure 1).

Then we assessed cardiac function in these mice mainly via echocardiography and LV catheterization. Lepr^db/db^ female mice have normal ejection fraction (≥50%) during at least their first year of life (Figure 1A). However, they display diastolic dysfunction attested by a significantly increased end diastolic pressure (EDP) from 3 months of age (Figure 1B). However, the maximum and minimum dP/dt, the relaxation time constant Tau and the heart rate were not modified at rest (Supplemental Figure 2 A-D). Increased EDP was associated with increased heart weight/tibia length ratio and increased diastolic left ventricular (LV) posterior wall thickness attesting cardiac hypertrophy (Figure 1C-D). Diastolic interventricular septum thickness and LV internal diameter were not different from that of control mice (Figure 1E-F). Consistent with cardiac hypertrophy, cardiomyocyte were significantly larger in 3 month old Lepr^db/db^ female mice (Figure 1G, H). Myocyte dedifferentiation, attested by Myosin-7 (Myh7) over expression appears later on at 6 month of age (Figure 1I). Notably, while both cardiac ANP and BNP mRNA expression increased with age, they were not different in Lepr^db/db^ mice and control mice (Supplemental Figure 2 E-F).

**Figure 1:**
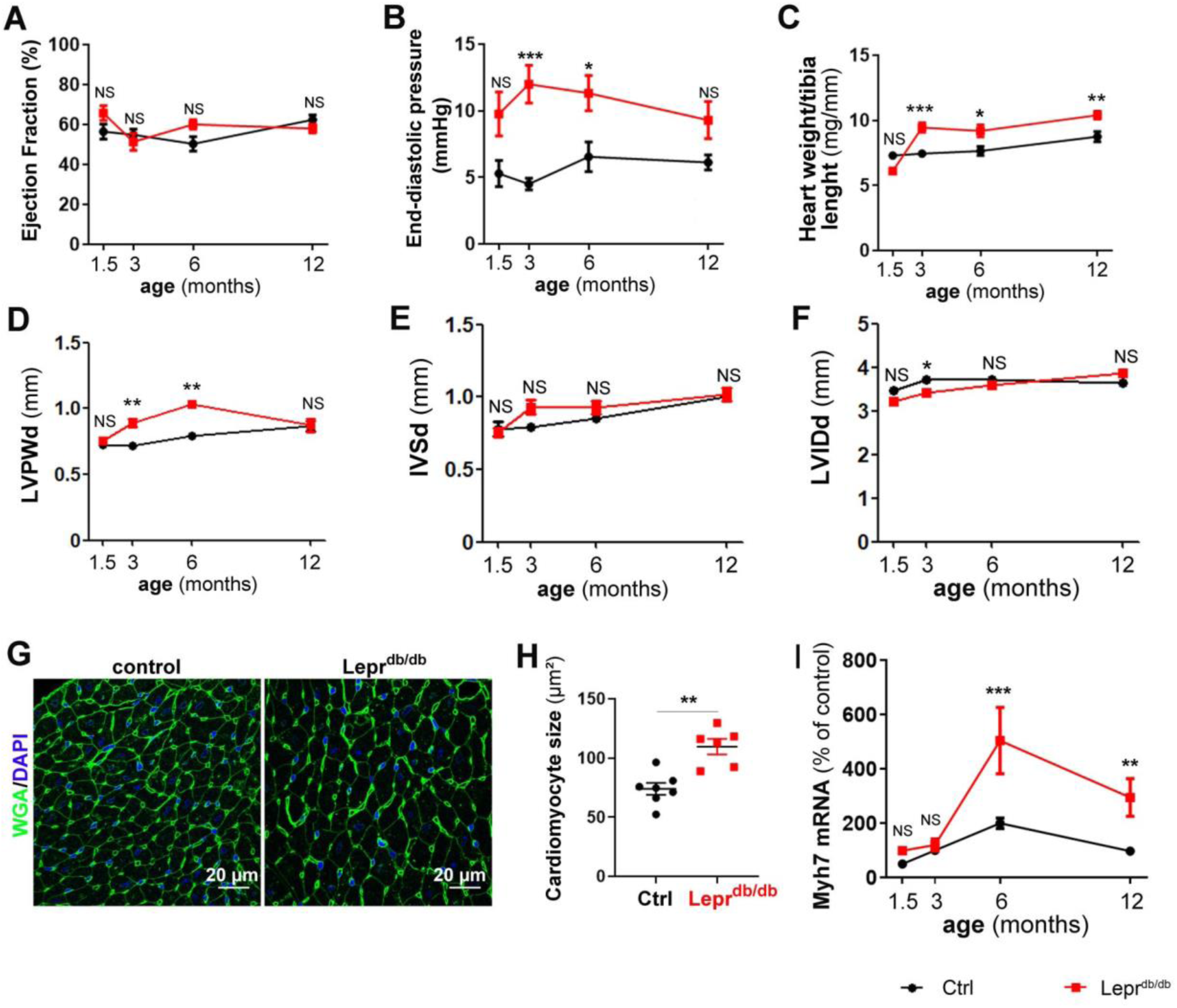
Lepr^db/db^ female mice have diastolic dysfunction from 3 months of age. Lepr^db/db^ female mice and their control Lepr^db^/+ littermates were subjected to echocardiography, LV catheterization and sacrificed at the indicated time points (n=8 to 15 per group). (**A**) Ejection faction was measured via echocardiography. (**B**) End diastolic pressure was measured using a pressure catheter. (**C**) Heart hypertrophy was assessed by calculating the heart weight over tibia length ratio. (**D**) End diastolic left ventricular posterior wall thickness during (LVPWd) was measured via echocardiography. (**E**) End diastolic Interventricular septum thickness (IVSd) was measured via echocardiography. (**F**) End diastolic Left ventricular internal diameter was measured via echocardiography. (**G**) Heart cross sections from 3 month old mice were stained with FITC-labeled WGA. Representative pictures are shown. (**H**) The mean cardiomyocyte surface area was measured. (**I**) Myh7 mRNA expression was measured via RT-qPCR in heart biopsies. *: p≤0.05; **: p≤0.01. ***: p≤0.001. NS: not significant. Two way ANOVA followed by Bonferroni’s multiple comparisons test or Mann Whitney test.

Altogether, this first set of data confirmed that Lepr^db/db^ mice have diastolic dysfunction with preserved systolic function and may be used as a mouse model of HFpEF ^13,14^.

### Lepr^db/db^ female mice do not have cardiomyocyte gross anomalies

To search for cardiomyocyte major anomalies, we first performed Desmin and sarcomeric alpha actin staining on heart cross sections to visualize cardiomyocyte sarcomeres and Connexin 43 (Cx43) to identify intercalary disks. We did not observe any significant difference between Lepr^db/db^ mice and their control littermates (Supplemental Figure 3A-B). Then we performed RT-qPCR to measure expression level of several cardiomyocyte markers notably, Titin isoforms N2-A and N2-B, Myocyte-specific enhancer factor 2C (MEF2C), Cx43, Sarcoplasmic/endoplasmic reticulum calcium ATPase 2 (Serca2) and Cardiac phospholamban (Pln) (Supplemental Figure 3C-H). None of these markers were significantly different in Lepr^db/db^ mice compared to their control littermates.

In conclusion, Lepr^db/db^ female mice do not have cardiomyocyte gross anomalies, which is consistent with the unchanged dP/dT minimum and maximum.

### Lepr^db/db^ female mice do not have significant cardiac fibrosis

Cardiac fibrosis, often associated with HFpEF, was first evaluated via picroSirius red staining. As shown in supplemental figure 3, the red staining was equivalent in 3 month old Lepr^db/db^ and in their control littermates (Supplemental Figure 4A-B). This result was confirmed using the Sircol™ collagen assay (Supplemental Figure 4C). Moreover neither collagen type I alpha 1 chain (Col1a1), nor collagen type III alpha 1 chain(Col3a1) mRNA expression was modified (Supplemental Figure 4D-E). Finally, no difference in Lysyl oxidase (Lox) mRNA expression, an enzyme promoting collagen cross linking was observed (Supplemental Figure 4F). To test whether cardiac fibrosis may appear later in life, we performed the same assays in a year old animals. Similarly to what we found in 3 month old animal, neither picrosirius red stain nor the Sircol™ collagen assay showed any differences between Lepr^db/db^ mice and control littermate (Supplemental Figure 4G-H). Although Lox mRNA was significantly increased in a year old Lepr^db/db^ mice compared to their control littermate (Supplemental Figure 4J), it did not lead to increased amount of insoluble collagen in the heart (Supplemental Figure 4K).

**Figure 2:**
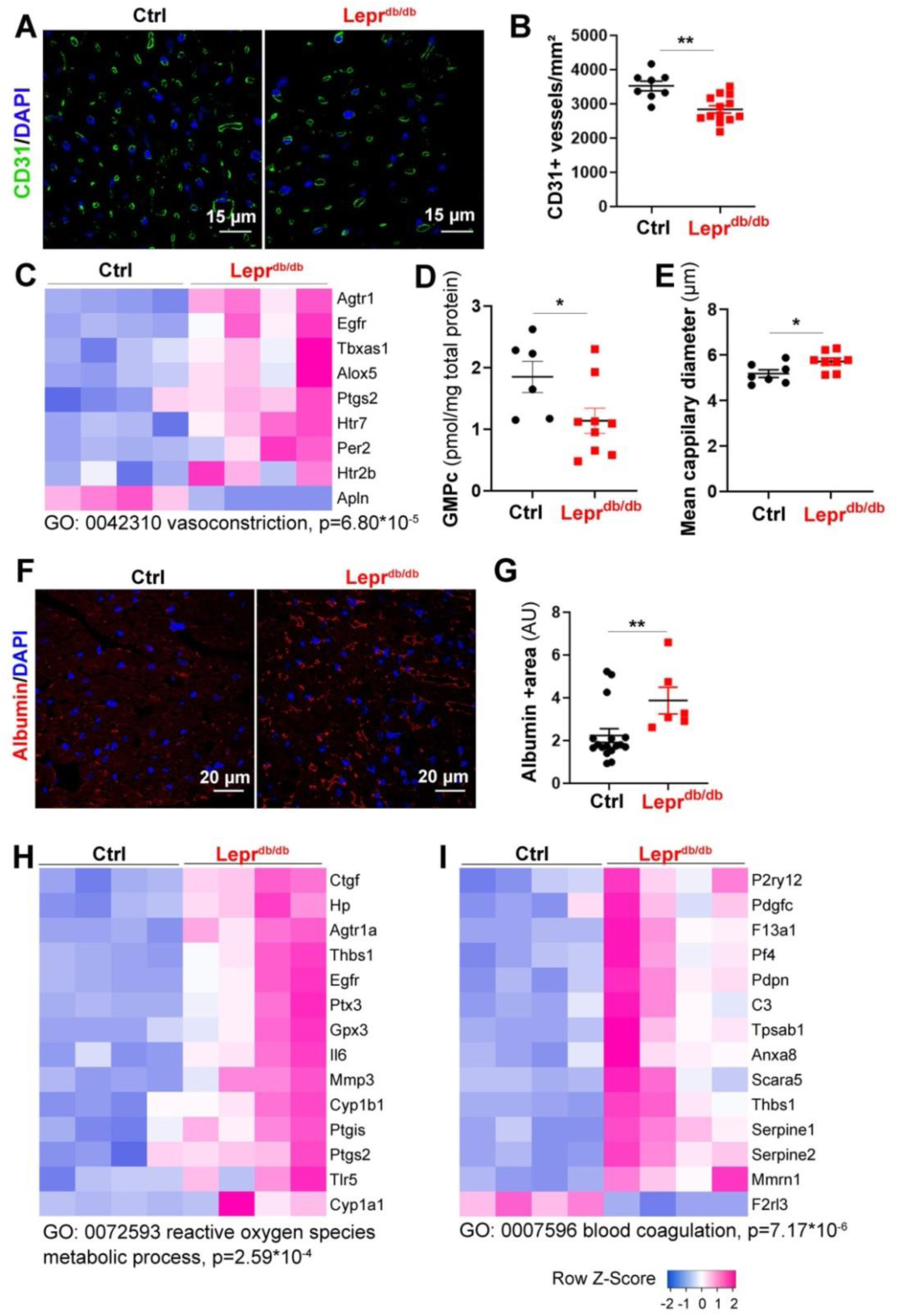
Lepr^db/db^ female mice have cardiac microvascular dysfunction. Lepr^db/db^ female mice and their control Lepr^db^/+ littermates were sacrificed at 3 months of age. (**A**) Heart cross sections were immune-stained with anti-CD31 antibodies to identify ECs. Representative pictures are shown. (**B**) Capillary density was quantified as the number of CD31+ vessels/mm^2^ (n=14 and 8 mice/ group). (**C**) RNA sequencing analysis of the cardiac vascular fraction revealed that “vasoconstriction” is one of the biological processes significantly increased in Lepr^db/db^ mice (n=4 mice/group). (**D**) The GMPc content of total aortas was quantified by ELISA (n=9 and 6 mice/group). (**E**) The mean cardiac capillary diameter was measured (n=7 and 6 mice/ group). (**F**) Heart cross sections were immuno-stained with anti-Albumin antibodies to assess vascular leakage. Representative pictures are shown. **(G)** Albumin extravasation was measured as the albumin+ surface area. ^2^ (n=6 and 15 mice/ group). **(H)** RNA sequencing analysis of the cardiac vascular fraction revealed that “Reactive oxygen species metabolic process” is one of the biological processes significantly increased in Lepr^db/db^ mice (n=4 mice/group). (**I**) RNA sequencing analysis of the cardiac vascular fraction revealed that “Blood coagulation” is one of the biological processes significantly increased in Lepr^db/db^ mice (n=4 mice/group). **: p≤0.01. Mann Whitney test.

**Figure 3:**
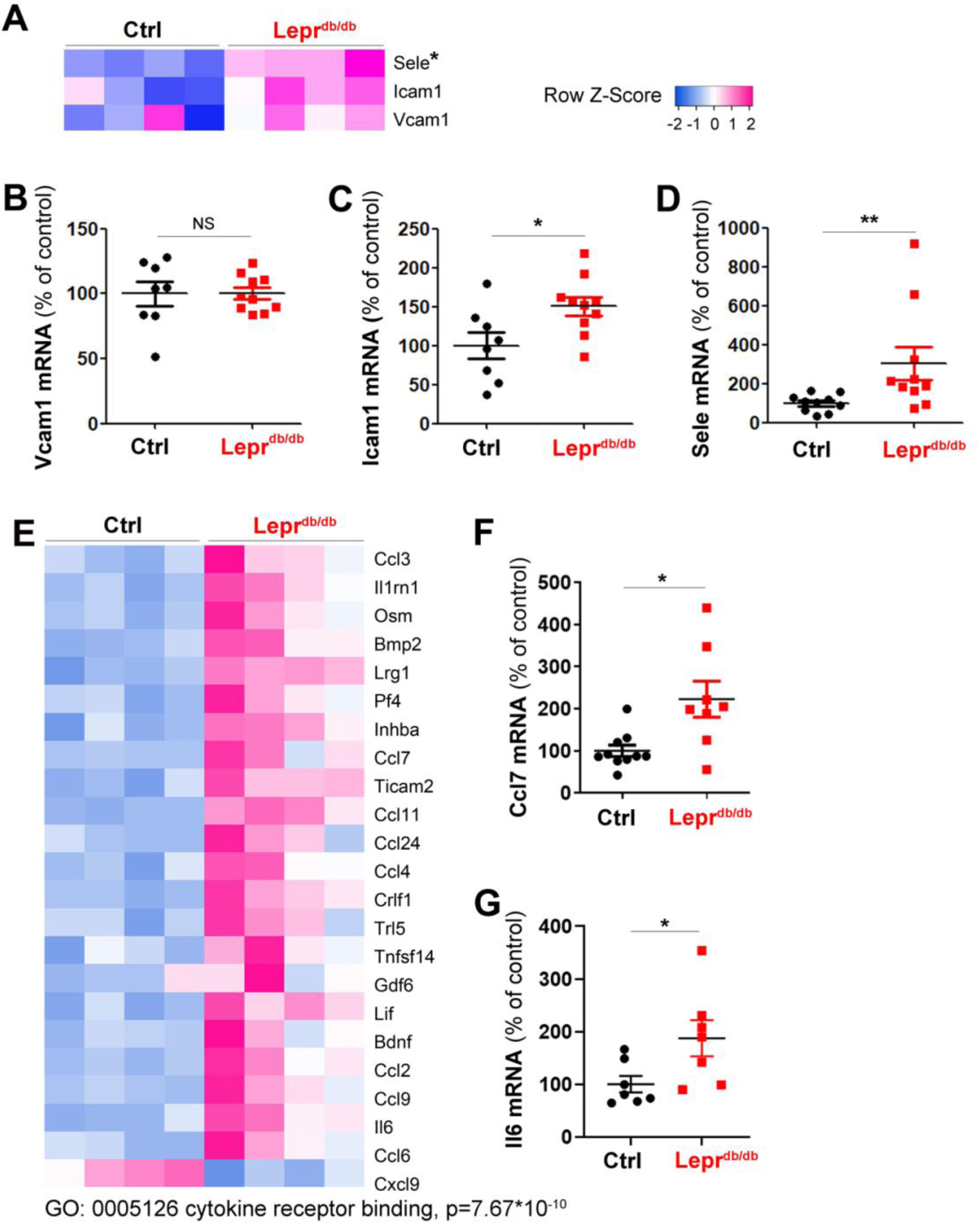
Lepr^db/db^ female mice have increased cardiac inflammation. Lepr^db/db^ female mice and their control Lepr^db^/+ littermates were sacrificed at 3 months of age. (**A**) RNA sequencing analysis of the cardiac vascular fraction revealed an increased expression of adhesion molecules. Vcam1 (**B**), Icam1 **(C)** and Sele (**D**) mRNA expression was measured via RT-qPCR in total heart biopsies (n=10 mice/group). RNA sequencing analysis of the cardiac vascular fraction revealed that “Cytokine receptor binding” is one of the molecular function significantly increased in Lepr^db/db^ mice (n=4 mice/group). Ccl7 (**F**) and Il-6 (**G**) mRNA expression was measured via RT-qPCR in total heart biopsies (n=10 mice/group). *: p≤0.05; **: p≤0.01. NS: not significant. Mann Whitney test.

In conclusion, diastolic dysfunction in Lepr^db/db^ female mice is not associated with increased cardiac fibrosis.

### 3 month old Lepr^db/db^ female mice have cardiac microvessel disease

To phenotype the cardiac microvessel of 3 month old Lepr^db/db^ female mice, we performed both immuno-histological and gene expression analyses. Gene expression analysis was performed by a transcriptomic analysis via RNA sequencing of the cardiac vascular fraction which includes ECs, pericytes, few smooth muscle cells and few perivascular inflammatory cells (Supplemental Table 2). The main RNA sequencing results were confirmed by RT-qPCR performed on total heart RNA samples.

First we quantified cardiac capillary density and found that capillary density is significantly decreased in the heart of Lepr^db/db^ female mice (Figure 2A-B) which is fully consistent with previous analyses performed on human patient with HFpEF ^15^. Transcriptomic analysis via RNA sequencing of the cardiac vascular fraction of these mice showed a switch in the balance between vasodilator and vasoconstrictor molecules in favor of vasoconstriction of cardiac arterioles (Figure 2C) notably, because of the downregulated expression of Apelin (Apln) and the increased expression of Type-1 angiotensin II receptor (Agtr1), Thromboxane-A synthase (Tbxas1), Arachidonate 5-lipoxygenase (Alox5) and Prostaglandin G/H synthase 2 (Ptgs2) (Figure 2C). This result is consistent with the decreased GMPc content of the aorta of Lepr^db/db^ mice indicating decreased guanylate cyclase activity (Figure2D). However, when we measured the mean cardiac capillary diameter (notably cardiac capillaries are not muscularized), we found that it was significantly increased in Lepr^db/db^ mice compared to control mice (Figure 2E). To assess vascular permeability we measure albumin extravasation and found a significantly increased albumin extravasation in Lepr^db/db^ mice compared to their control littermate which indicate abnormal vessel permeability (Figure 2F, G). Besides, the transcriptomic analysis of the cardiac vascular fraction also revealed oxidative stress attested by Cytochrome P450 1B1 (Cyp1b1) and Cytochrome P450 1A1 (Cyp1a1) upregulation (Figure 2H) and a pro-coagulation phenotype attested by P2Y purinoceptor 12 (P2ry12), Coagulation factor XIII A chain (F13a1) and Platelet factor 4 (PF4) upregulation (Figure 2I). Finally, both the transcriptomic analysis of the cardiac vascular fraction and gene expression analysis performed on total heart extract showed a significantly increased expression of adhesion molecule Icam1 and E-selectin (Figure 3A-D) demonstrating EC activation. EC activation was associated with upregulation of multiple inflammatory cytokines including Interleukin-6 (Il6), C-C motif chemokine 2 (Ccl2), C-C motif chemokine 3 (Ccl3) and C-C motif chemokine 7 (Ccl7) (Figure 3E-G).

Altogether, these results demonstrate that Lepr^db/db^ have cardiac microvessel disease characterized by a decreased capillary density but an increased capillary diameter. Moreover, they display oxidative stress together with a pro-inflammatory and a pro-coagulant phenotype.

### 3 month old Lepr^db/db^ female mice have cardiac inflammation

In association with cardiac EC activation, leucocyte infiltration in the cardiac tissue of Ler^db/db^ mice was significantly increased (Figure 4A-B). Among Leukocytes, cells were mostly CD68+/Mrc1+ M2 macrophages (Supplemental Figure 5A-B). Besides, leukocytes also included CD3+ T-lymphocytes (Supplemental Figure 5C-D), Ly6G+ neutrophils (Supplemental Figure 5E-F) and B220 B-lymphocytes (Supplemental Figure 5G-H). Both M2 macrophage and B-lymphocyte densities were increased in Lepr^db/db^ mice. However, T-lymphocytes and neutrophils densities were not different between Lepr^db/db^ mice and their control littermates.

**Figure 4:**
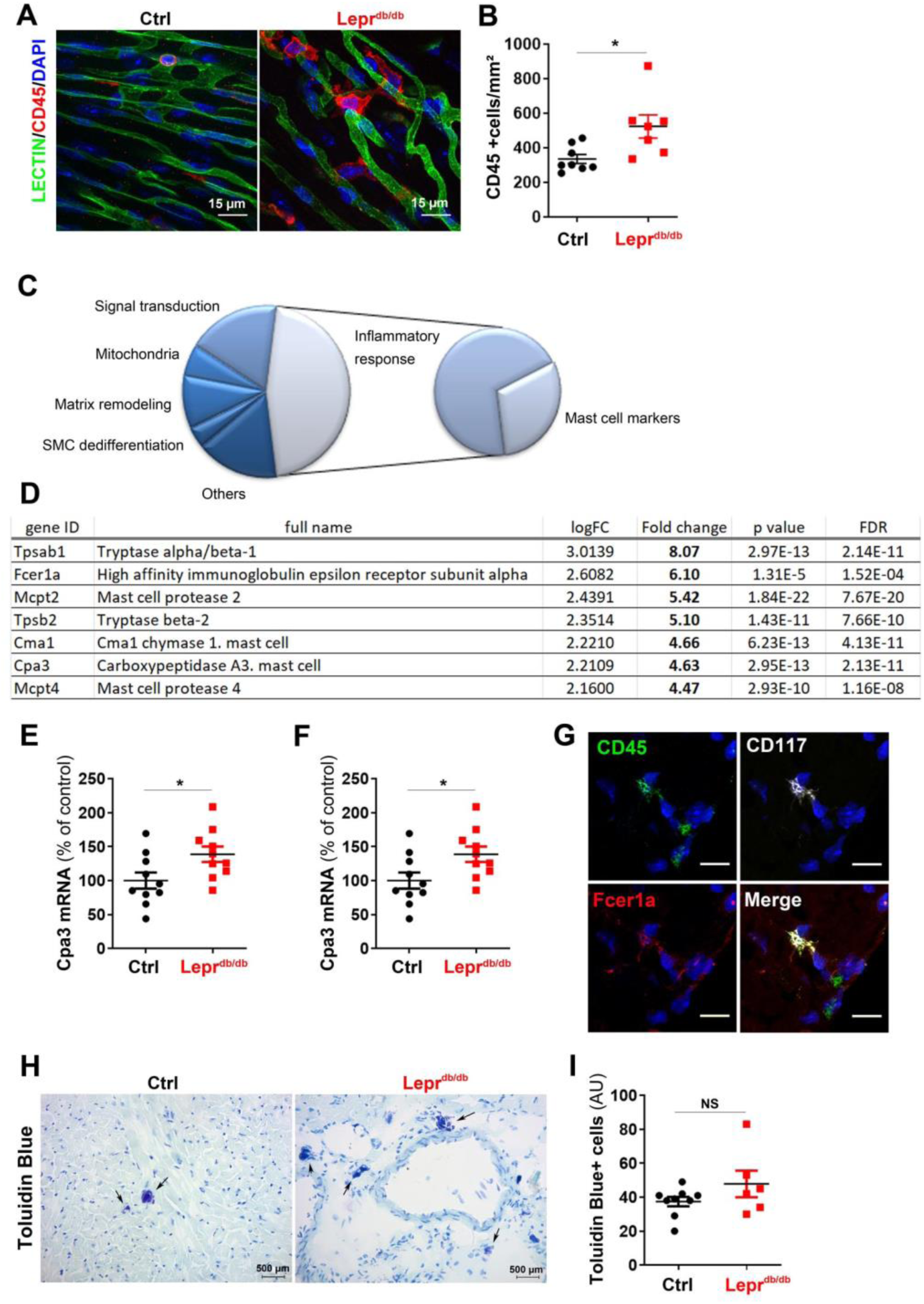
Lepr^db/db^ female mice have increased cardiac inflammation. Lepr^db/db^ female mice and their control Lepr^db^/+ littermates were sacrificed at 3 months of age. (**A**) BS1-lectin-FITC-perfused heart cross sections were immuno-stained with anti-CD45 antibodies to assess cardiac inflammation. Representative pictures are shown. (**B**) Leucocyte infiltration was measured as the number of CD45+ cells/mm^2^ (n=7 and 8 mice/ group). (**C**) RNA sequencing analysis of the cardiac vascular fraction was performed. Among the 50 most upregulated gene in Lepr^db/db^ female mice, 7 are mast cell markers. **(D)** List of mast cell marker upregulated in Lepr^db/db^ female mice. Cpa3 (**E**) and Fcer1a (**F**) mRNA expression was measured via RT-qPCR in total heart biopsies (n=10 mice/group). (**G**) Heart cross sections were co-immunostained with anti-CD45 (in green), anti-CD117 (in white) and anti-Fcer1a (in red) antibodies. (**H**) Heart cross sections were stained with Toluidine Blue to identify mast cells. Representative pictures are shown. (**I**) Mast cell infiltration was quantified as the number of toluidine blue+ cells/heart section. *: p≤0.05; NS: not significant. Mann Whitney test.

### 3 month old Lepr^db/db^ female mice have abnormal cardiac mast cell activation

Apart from highlighting EC dysfunction and cardiac inflammation, transcriptomic analysis of the cardiac vascular fraction also revealed that among the 50 genes with log2FC≥2, 7 genes were mast cell markers (Figure 4C-D). We confirmed Carboxypeptidase A3, mast cell (Cpa3) and High affinity immunoglobulin epsilon receptor subunit alpha (Fcer1a) overexpression in total heart extracts (Figure 4E-F) and confirmed that Fcer1a was indeed expressed by CD45+, CD117+ cardiac mast cell (Figure 4G). We then quantified the total number of cardiac mast cell after Toluidine blue staining of heart section but found no significant differences between Lepr^db/db^ mice and their control littermate however (Figure 4H-I), when we calculated the percentage of activated/degranulating mast cells, we found that it was significantly increased in the heart of Lepr^db/db^ mice (Figure 5A-B). Since mast cells are typically activated by IgE, we measured IgE circulating levels in plasma extracts from Lepr^db/db^ mice and control littermates. Consistent with increased mast cell activated IgE circulating level were significantly increased in Lepr^db/db^ mice (Figure 5C-D).

**Figure 5:**
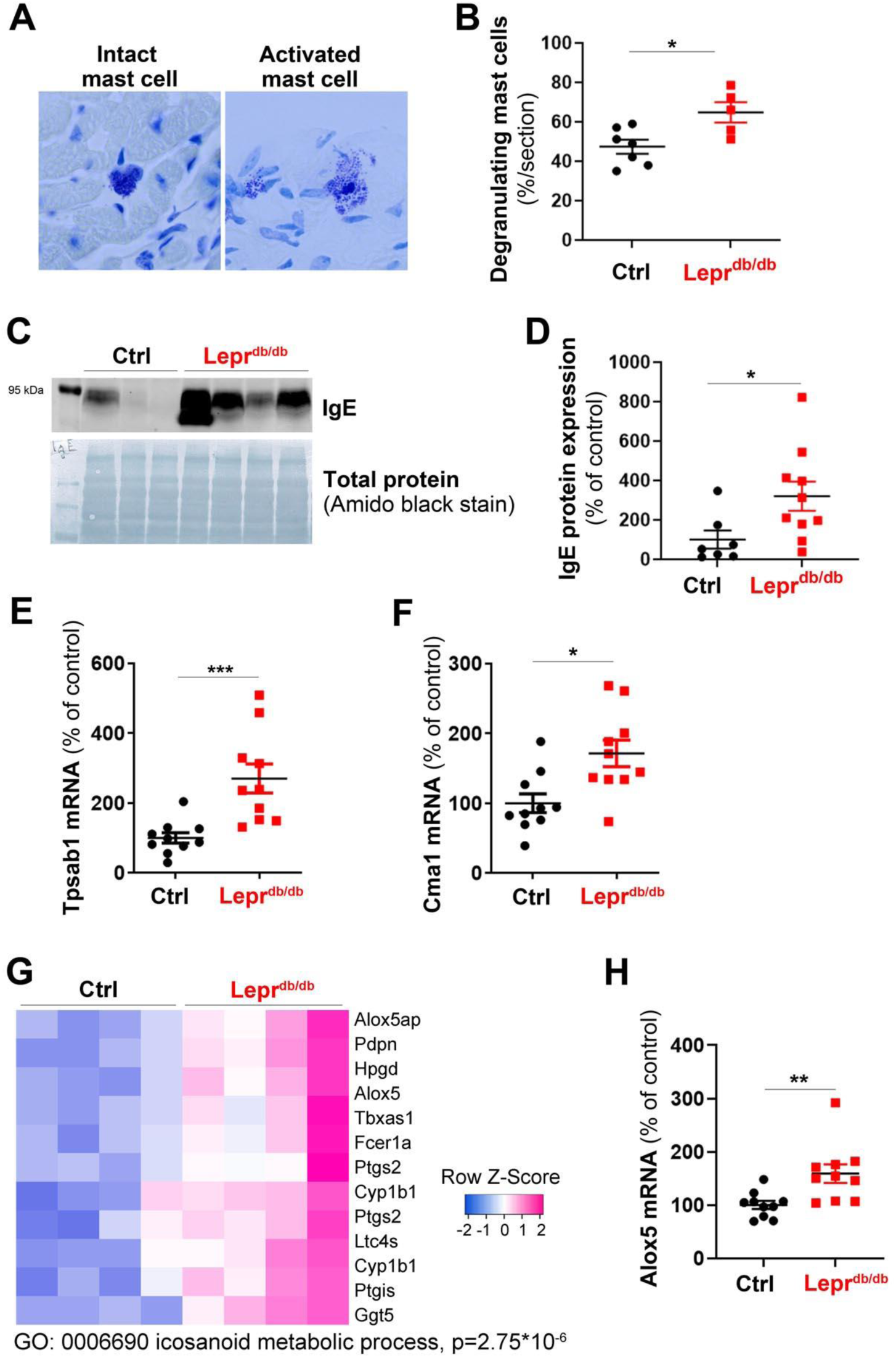
Lepr^db/db^ female mice have increased circulating IgE and cardiac mast cell activation. Lepr^db/db^ female mice and their control Lepr^db^/+ littermates were sacrificed at 3 months of age. (**A**) Heart cross sections were stained with Toluidine Blue to identify mast cells degranulation. (**B**) The percentage of degranulating mast cell was quantified. (**C**) Plasmatic IgE level was assessed by western blot analysis and (**D**) quantified (n=10 and 7 mice/group). Tpsab1 (**E**) and Cma1 (**F**) mRNA expression was measured via RT-qPCR in total heart biopsies (n=10 mice/group). (**G**) RNA sequencing analysis of the cardiac vascular fraction revealed that “Icosanoid metabolic process” is one of the molecular function significantly increased in Lepr^db/db^ mice (n=4 mice/group). (**H**) Alox5 mRNA expression was measured via RT-qPCR in total heart biopsies (n=10 mice/group). *: p≤0.05; **: p≤0.01; ***: p≤0.001. Mann Whitney test.

Mast cells are known to induce cardiovascular effects through the release of their granule content ^16^, notably proteases, histamine and the production of eicosanoids and cytokines. Consistent with increased cardiac mast cell activation, both Tryptase (Tpsab1) and Chymase (Cma1) mRNA were significantly increased in the heart of Lepr^db/db^ mice compared to the one of their control littermates (Figure 5E-F). Also the transcriptomic analysis of the cardiac vascular fraction revealed increased eicosanoid metabolic process attested by increased arachidonate 5-lipoxygenase (Alox5), thromboxane A synthase 1 (Tbxas1) and leukotriene C4 synthase (Ltc4s) expression (Figure 5G). The increased Alox5 mRNA expression was confirmed in total heart samples (Figure 5H).

### Activated cardiac mast cells promote the appearance of diastolic dysfunction in Lepr^db/db^ mice

To investigate the role of mast cell degranulation in the pathophysiology of diastolic dysfunction in Lepr^db/db^ mice, 2 month old Lepr^db/db^ mice (i.e. before they display increased EDP) were treated with 50 mg/Kg/day cromolyn sodium versus vehicle. Mice were sacrificed 28 days later. First, we verified cromolyn sodium therapy was effective and did decrease the percentage of degranulating mast cells in the heart (Figure 6A), and then we compared cardiac function in cromolyn sodium treated and vehicle-treated Lepr^db/db^ mice. Ejection fraction, which is normal in Lepr^db/db^ mice was not modulated (Figure 6B), however, EDP was significantly decreased in cromolyn sodium treated Lepr^db/db^ mice (Figure 6C) indicating improved diastolic function. Cromolyn sodium therapy did neither modify the heart weight nor the LV posterior wall thickness nor the mean cardiomyocyte size (Figure 6 D-G), indicating that cromolyn sodium therapy does not prevent cardiac hypertrophy.

**Figure 6:**
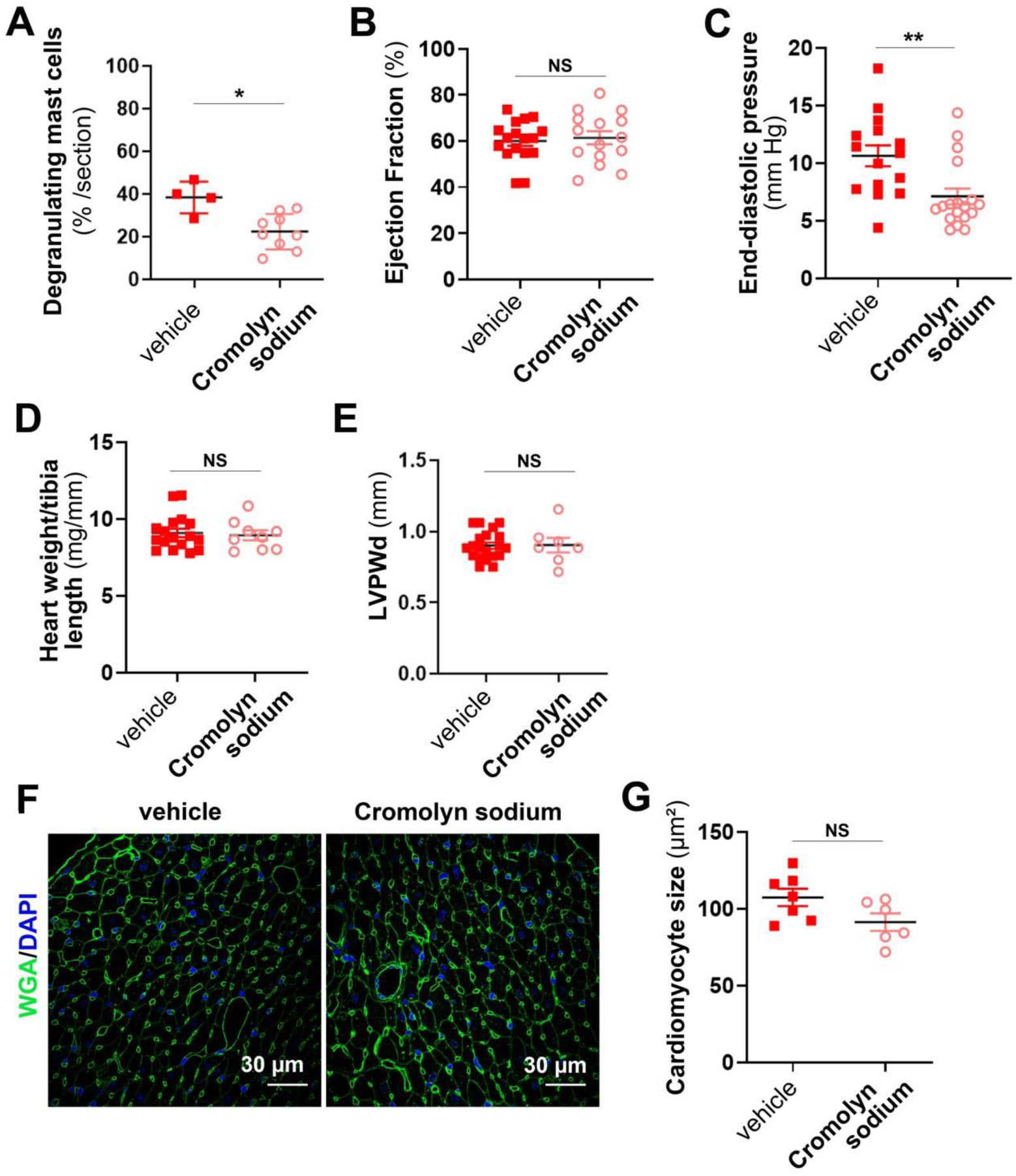
Cromolyn sodium improves diastolic function in Lepr^db/db^ female mice. 2 month old Lepr^db/db^ female mice were treated or not with 50 mg/kg/day cromolyn sodium for 28 days. At 3 month of age. Mice were subjected to echocardiography, LV catheterization and sacrificed (**A**) Heart cross sections were stained with Toluidine Blue to identify mast cells. Mast cell activation was quantified as the percentage of degranulating mast cells. (**B**) Ejection faction was measured via echocardiography. (**C**) End diastolic pressure was measured using a pressure catheter. (**D**) Heart hypertrophy was assessed by calculating the heart weight over tibia length ratio. (**E**) End diastolic left ventricular posterior wall thickness during (LVPWd) was measured via echocardiography. (**F**) Heart cross sections were stained with FITC-labeled WGA. Representative pictures are shown. (**H**) The mean cardiomyocyte surface area was measured. *: p≤0.05; **: p≤0.01; NS: not significant. Mann Whitney test.

Altogether these results indicated that mast cells, via secretion of their granule content, promote the development of diastolic dysfunction, however, they do not participate in the development of cardiomyocyte hypertrophy.

To investigate mechanism by which mast cell may induce diastolic dysfunction we investigated the effect of cromolyn sodium therapy on cardiac microvessel phenotype and cardiac inflammation. First we found that cromolyn sodium therapy does not modify cardiac microvessel density (Figure 7A-B), however it did reduce capillary diameter (Figure 7C) and permeability attested by decreased albumin extravasation (Figure 7D-E). Moreover, CD45+ leucocyte recruitment was significantly decreased (Figure 7F-G).

**Figure 7:**
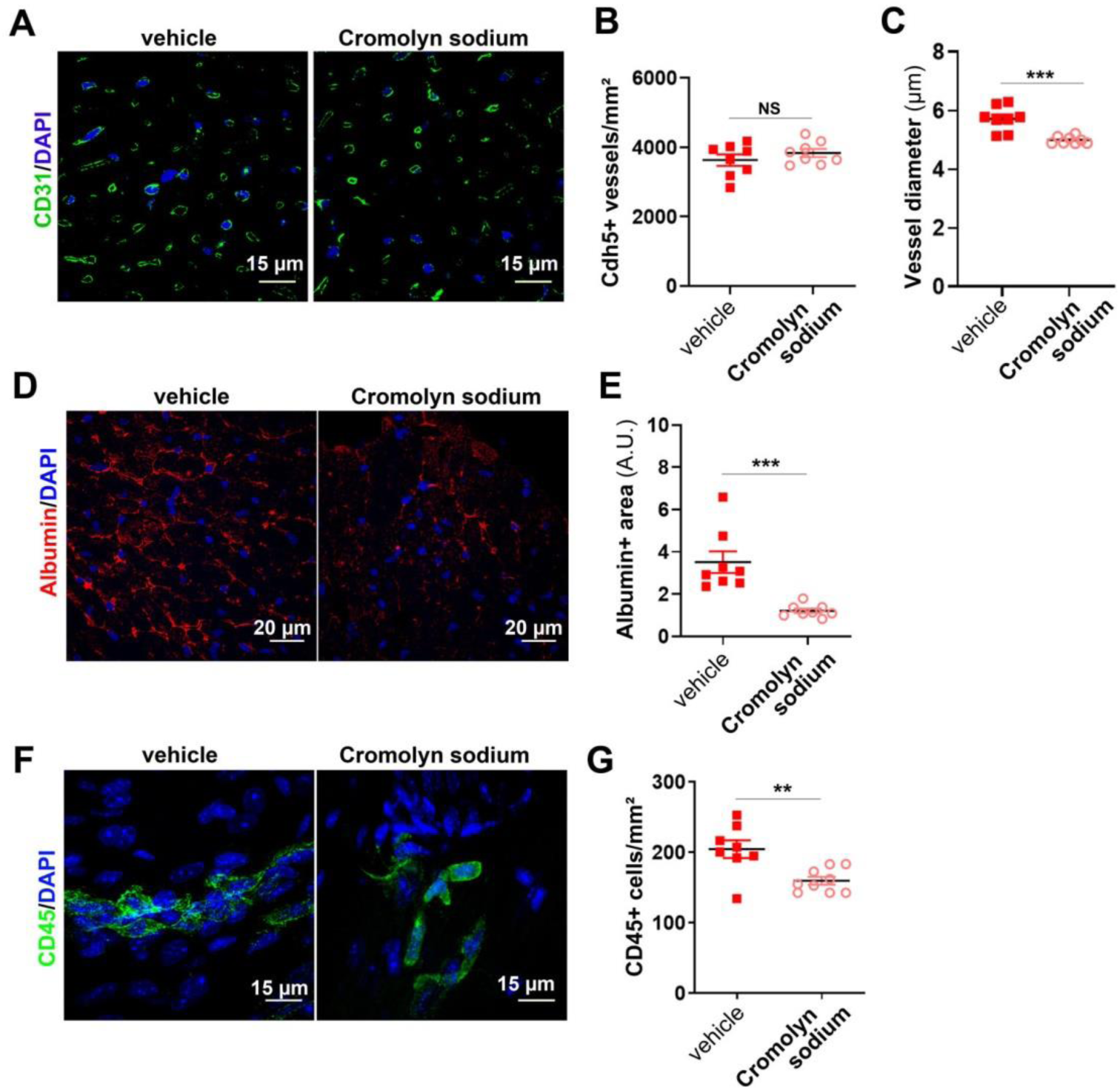
Cromolyn sodium decreases vascular permeability and leucocyte infiltration in Lepr^db/db^ female mice. 2 month old Lepr^db/db^ female mice were treated or not with 50 mg/kg/day cromolyn sodium for 28 days. At 3 month of age. Mice were subjected to echocardiography, LV catheterization and sacrificed. (**A**) Heart cross sections were immune-stained with anti-CD31 antibodies to identify ECs. Representative pictures are shown. (**B**) Capillary density was quantified as the number of CD31+ vessels/mm^2^ (n= 8 mice/ group). (**C**) The mean cardiac capillary diameter was measured (n= 8 mice/ group). (**D**) Heart cross sections were immunostained with anti-albumin antibodies to assess vascular leakage. Representative pictures are shown. (**E**) Albumin extravasation was measured as the albumin+ surface area (n=8 mice/ group). (**F**) Heart cross sections were immunostained with anti-CD45 antibodies to assess cardiac inflammation. Representative pictures are shown. (**G**) Leucocyte infiltration was measured as the number of CD45+ cells/mm^2^ (n=10 mice/ group). *: p≤0.05; **: p≤0.01; ***: p≤0.001. NS: not significant. Mann Whitney test.

In conclusion, mast cells promote cardiac capillary permeability and vasodilation, which is consistent with the well-known effect of histamine contained in mast cell granules ^17^. Moreover, mast cells participate in cardiac leucocyte infiltration.

## Discussion

The present study supports the microvascular hypothesis of HFpEF especially in the setting of obesity and type 2 diabetes. In this paper, we used Lepr^db/db^ female mice as a model of diastolic dysfunction. Lepr^db/db^ female mice have the advantage of recapitulating the main risk factors for HFpEF, i.e. diabetes, obesity female gender and hypertension. Lepr^db/db^ mice were previously shown to display diastolic dysfunction ^13,18,19^ and to recapitulate significant features of human HFpEF ^14,20^. In the present study, we thoroughly characterized the cardiac microvascular phenotype of these mice, notably, via a transcriptomic analysis. Notably we revealed that cardiac microvessel disease is characterized by a decreased capillary density, abnormal vessel permeability and vasoconstriction of arterioles but increased capillary diameter; moreover we showed that ECs display oxidative stress and have a pro-inflammatory and pro-coagulant phenotype. Strikingly, we demonstrated for the first time that, in Lepr^db/db^ mice, cardiac microvessel disease is associated with increased mast cell activation and proved that it participates to the pathophysiology of both cardiac microvessel disease and diastolic dysfunction (Supplemental Figure 6).

The current paradigm for HFpEF proposes that myocardial remodelling and dysfunction in HFpEF results from the following sequence of events: 1) comorbidities including obesity, diabetes and/or hypertension would induce a systemic low grade pro-inflammatory state; 2) this pro-inflammatory state would induce EC dysfunction characterized by an increased ROS production, a decreased NO synthesis and an increased expression of adhesion molecules such as VCAM-1 and E-selectin; 3) EC dysfunction would lead to a compromised heart perfusion secondary to impaired NO-dependent vasodilatation, oedema and pro-inflammatory/pro-thrombotic phenotype, macrophage infiltration and fibrosis ^4,5^. The present data show that the Lepr^db/db^ female mice model largely recapitulates this paradigm while adding further features. Notably, we demonstrated that mast cell activation, which is either part of the low grade pro-inflammatory state or induced by the low grade inflammatory state of diabetic obese mice, promotes microvascular dysfunction especially vascular permeability and capillary dilation and participates in the development of diastolic dysfunction. However, EC activation may precede mast cell activation (data not shown). Consistent with a central role of inflammation ^21^ and microvascular disease in the pathophysiology of HFpEF, Lepr^db/db^ mice do not display significant cardiac fibrosis or major cardiomyocyte abnormalities. Notably, we found that cardiomyocyte hypertrophy does not seem to promote the increased EDP. Indeed, although cromolyn sodium therapy prevents increased EDP, this effect is not associated to cardiomyocyte hypertrophy and dedifferentiation after the onset of diastolic dysfunction, as demonstrated by Myh7 overexpression.

Mast cells are immune cells that reside in the connective tissues including the myocardium. They are characterized by the expression of c-Kit receptors and by their granules containing active mediators including proteases, notably Cma1, Tpsab1 and histamine. Mast cells may be activated by IgEs via their receptor Fcer1a, Complement factors via Toll-like receptors, IgGs or cytokines ^22^. They have been associated with several cardiovascular diseases including atherosclerosis, myocardial infarction and aneurysms ^16^, pathologies in which mast cells are contributing to the pathogenesis essentially through the release of their granule content. Importantly, circulating Tryptase was recently suggested to be a marker for cardiovascular diseases ^23^. Moreover, mast cells have been previously involved in diastolic dysfunction induced by ovariectomy in rats ^24^ and diabetic cardiomyopathy in streptozotocin-treated mice ^25^. The present study thus confirms the significant role of mast cells in cardiovascular diseases. How mast cells are activated in the setting of cardiovascular diseases remains unknown ^22^. We found that, in Lepr^db/db^ mice, increased activation of mast cells is associated with increased circulating levels of IgEs. IgE/Fcer1a is the main route of mast cell activation in allergic diseases. Interestingly, IgEs were reported to be elevated in the serum of patients with cardiovascular diseases ^26 27 28^ including coronary arterial disease and myocardial infarction. Consistently, Asthma was shown to be related to an increased incidence of coronary heart disease, particularly in women ^29^. More specifically, one study reported that coronary flow reserve, considered as an early marker of endothelial dysfunction is significantly lower in patients with high IgE levels ^30^. Altogether, these results further support that Lepr^db/db^ mice are a relevant model of human cardiovascular diseases.

In conclusion, the present study further confirms that inflammation and cardiac microvessel disease are at the heart of HFpEF pathophysiology ^21^ and identified for the first time mast cells as critical players of cardiac microvessel disease and diastolic dysfunction, making them a promising therapeutic target for HFpEF treatment.

## Author contributions

S.G. conducted experiments, acquired data, analyzed data. L.C., P-L.H., C.C. and M-L.B. conducted experiments, acquired data. A.-P. G. critically revised the manuscript. T.C. designed research studies and critically revised the manuscript. M.-A. R. designed research studies, conducted experiments, acquired data, analyzed data and wrote the manuscript

## Acknowledgments

We thank Philippe Alzieu, Annabel Reynaud, Sylvain Grolleau, and Maxime David for their technical help. We thank Christelle Boullé for administrative assistance.

This work benefited from equipment and services from the iGenSeq (RNA sequencing) and iCONICS (RNAseq analysis) core facilities at the ICM (Institut du Cerveau et de la Moelle épinière, Hôpital Pitié-Salpêtrière, PARIS, France).

This study was supported by grants from the Fondation pour la Recherche Médicale (équipe FRM), from the Region « Nouvelle Aquitaine » and the Agence Nationale pour la Recherche (Appel à Projet Générique). Finally, this study was co-funded by the “Institut National de la Santé et de la Recherche Médicale” and by the University of Bordeaux.

## Conflicts of interest

none

## Notes

The authors have declared that no conflict of interest exists.

### Competing Interest Statement

The authors have declared no competing interest.

