## Supplemental Table 2 for "Mast cells participate in the development of diastolic dysfunction in diabetic obese mice"

| | FPKM | | | | | | | | | | | |
|---|---|---|---|---|---|---|---|---|---|---|---|---|
| FeatureID | Leprdb/+_1 | Leprdb/+_2 | Leprdb/+_3 | Leprdb/+_4 | Leprdb/db_1 | Leprdb/db_2 | Leprdb/db_3 | Leprdb/db_4 | logFC | PValue | FDR | Status |
| Chil3 | 0 | 0.03 | 0.03 | 0 | 0.89 | 1.08 | 1.07 | 2.71 | 6.098574681 | 1.08E-20 | 3.42E-18 | Up in group#2 |
| Cyp1a1 | 0.08 | 0.15 | 0.04 | 0.26 | 0.49 | 14.15 | 5.94 | 4.21 | 5.591431982 | 4.22E-14 | 3.60E-12 | Up in group#2 |
| Lvrn | 0.05 | 0 | 0.02 | 0.02 | 0.08 | 0.19 | 0.42 | 0.45 | 3.790473808 | 2.04E-08 | 5.08E-07 | Up in group#2 |
| Hif3a | 0.01 | 0.02 | 0.03 | 0.04 | 0.17 | 0.34 | 0.33 | 0.21 | 3.535038239 | 1.69E-16 | 2.21E-14 | Up in group#2 |
| Zbtb16 | 1.29 | 0.84 | 1.2 | 1.62 | 3.1 | 25.85 | 10.9 | 10.51 | 3.411311524 | 1.46E-15 | 1.61E-13 | Up in group#2 |
| Olfm2 | 0.12 | 0.03 | 0.17 | 0.24 | 1.15 | 1.19 | 1.94 | 1.32 | 3.38208536 | 2.34E-20 | 6.64E-18 | Up in group#2 |
| Elmod1 | 0.02 | 0.02 | 0.09 | 0.04 | 0.32 | 0.71 | 0.37 | 0.44 | 3.375994241 | 1.83E-11 | 9.54E-10 | Up in group#2 |
| Fcna | 1.02 | 0.98 | 2.19 | 1.5 | 6.67 | 7.51 | 24.15 | 12.65 | 3.211669671 | 1.76E-18 | 3.55E-16 | Up in group#2 |
| Gldc | 0.13 | 0.06 | 0.06 | 0.03 | 0.28 | 1.02 | 0.71 | 0.53 | 3.167697506 | 2.53E-12 | 1.57E-10 | Up in group#2 |
| Tpsab1 | 0.32 | 0.32 | 0.86 | 0.3 | 2.34 | 2.04 | 6.32 | 3.02 | 3.013997006 | 2.97E-13 | 2.14E-11 | Up in group#2 |
| Spock2 | 3.29 | 3.53 | 3.18 | 4.1 | 16.69 | 35.22 | 28.17 | 25.19 | 2.955923218 | 2.65E-60 | 3.53E-56 | Up in group#2 |
| Ptx3 | 0.89 | 0.56 | 0.69 | 0.54 | 2.32 | 2.96 | 7.6 | 6.07 | 2.867600718 | 3.25E-17 | 4.92E-15 | Up in group#2 |
| Spp1 | 0.51 | 0.64 | 0.33 | 0.32 | 2.33 | 3.29 | 4.31 | 2.87 | 2.862681922 | 4.14E-23 | 1.84E-20 | Up in group#2 |
| Has1 | 1.39 | 1.12 | 1.27 | 0.75 | 3.29 | 4.8 | 13.28 | 9.3 | 2.80347696 | 6.96E-16 | 8.00E-14 | Up in group#2 |
| Ugt1a6a | 0.03 | 0.03 | 0.14 | 0 | 0.23 | 0.21 | 0.46 | 0.5 | 2.778874077 | 1.81E-07 | 3.55E-06 | Up in group#2 |
| H2-T9 | 0 | 1.44 | 2.07 | 0 | 9.79 | 2.35 | 4.8 | 5.64 | 2.711314637 | 0.001961239 | 0.010331415 | Up in group#2 |
| Mrgprb1 | 0.04 | 0.08 | | 0.15 | 0.39 | 0.34 | 1.14 | 0.62 | 2.705625302 | 3.57E-10 | 1.38E-08 | Up in group#2 |
| Serpina3n | 1.77 | 0.9 | 1.45 | 0.73 | 4.99 | 7.28 | 8.47 | 9.6 | 2.691266369 | 7.75E-28 | 5.73E-25 | Up in group#2 |
| Pth1r | 3.23 | 3.18 | 2.9 | 4.17 | 14.16 | 27.29 | 23.38 | 18.27 | 2.681276152 | 1.25E-47 | 3.42E-44 | Up in group#2 |
| Coch | 0.08 | 0.16 | 0.36 | 0.28 | 1.35 | 1.63 | 1.37 | 0.95 | 2.627316214 | 4.07E-17 | 5.95E-15 | Up in group#2 |
| Fcer1a | 0 | 0.28 | 0.55 | 0.12 | 0.92 | 0.8 | 3.11 | 0.92 | 2.608297032 | 1.31E-05 | 0.000152092 | Up in group#2 |
| Hhipl1 | 0.2 | 0.32 | 0.13 | 0.31 | 0.79 | 2.12 | 1.4 | 1.13 | 2.545711298 | 3.77E-13 | 2.66E-11 | Up in group#2 |
| Ifi205 | 0.57 | 0.68 | 0.55 | 0.63 | 1.07 | 2.72 | 4.36 | 5.08 | 2.494049197 | 2.72E-11 | 1.38E-09 | Up in group#2 |
| Lcn2 | 1.09 | 0.77 | 1.31 | 1.66 | 6.02 | 6.18 | 7.85 | 6.17 | 2.492024048 | 1.47E-26 | 9.31E-24 | Up in group#2 |
| Mcpt2 | 3.39 | 4.85 | 5.75 | 6.64 | 19.72 | 18.41 | 43.21 | 26.45 | 2.439181977 | 1.84E-22 | 7.67E-20 | Up in group#2 |
| Fam84a | 0.22 | 0.36 | 0.36 | 0.28 | 1.3 | 1.41 | 2.49 | 1.19 | 2.422397153 | 3.28E-18 | 6.24E-16 | Up in group#2 |
| Pcdh20 | 0.03 | 0.04 | 0.1 | 0.07 | 0.22 | 0.32 | 0.39 | 0.29 | 2.4122122 | 7.96E-11 | 3.59E-09 | Up in group#2 |
| Tpsb2 | 1.13 | 0.98 | 2.14 | 2.27 | 4.06 | 5.81 | 14.1 | 8.01 | 2.351439975 | 1.44E-11 | 7.67E-10 | Up in group#2 |
| Vsig4 | 0.54 | 1.41 | 3.79 | 2.92 | 7.12 | 6.21 | 16.56 | 11.1 | 2.29768423 | 1.65E-08 | 4.20E-07 | Up in group#2 |
| Trem2 | 0.56 | 0.3 | 0.96 | 0.88 | 2.41 | 2.71 | 5.67 | 2.02 | 2.28449574 | 1.28E-09 | 4.33E-08 | Up in group#2 |
| Doc2b | 0.11 | 0.15 | 0.07 | 0.08 | 0.21 | 0.84 | 0.5 | 0.39 | 2.27523041 | 2.23E-08 | 5.46E-07 | Up in group#2 |
| Gpnmb | 0.17 | 0.18 | 0.47 | 0.18 | 0.82 | 1.1 | 1.35 | 1.3 | 2.230084948 | 2.07E-14 | 1.90E-12 | Up in group#2 |
| Tfec | 0.09 | 0.06 | 0.23 | 0.09 | 0.3 | 0.47 | 0.93 | 0.48 | 2.22340147 | 6.94E-06 | 8.70E-05 | Up in group#2 |
| Cma1 | 2.3 | 2.26 | 5.95 | 3.93 | 12.67 | 9.6 | 26.43 | 16.18 | 2.221091549 | 6.24E-13 | 4.13E-11 | Up in group#2 |
| Crlf1 | 0.23 | 0.12 | 0.25 | 0.23 | 0.81 | 0.68 | 1.37 | 0.95 | 2.21778152 | 2.40E-09 | 7.46E-08 | Up in group#2 |
| Cpa3 | 1.71 | 2.23 | 2.93 | 3.22 | 7.91 | 6.04 | 20.2 | 10.98 | 2.210989269 | 2.95E-13 | 2.14E-11 | Up in group#2 |
| Ccl7 | 5 | 5.01 | 6.55 | 5.5 | 10.11 | 18.56 | 38.51 | 30.8 | 2.208199933 | 2.67E-13 | 1.95E-11 | Up in group#2 |
| Fkbp5 | 4.51 | 2.91 | 2.98 | 4.24 | 9 | 23.51 | 15.82 | 16.04 | 2.193327407 | 4.28E-22 | 1.63E-19 | Up in group#2 |
| Cd209b | 0.14 | 0.08 | 0.16 | 0.17 | 0.25 | 0.28 | 1.42 | 0.39 | 2.191731139 | 0.000148496 | 0.001213944 | Up in group#2 |
| Mmp3 | 0.57 | 0.92 | 0.51 | 0.68 | 1.48 | 3.26 | 3.82 | 3.17 | 2.183348284 | 1.15E-13 | 8.89E-12 | Up in group#2 |

| | | | | | | | | | | | | |
|---|---|---|---|---|---|---|---|---|---|---|---|---|
| Mcpt4 | 2.71 | 3.18 | 7.9 | 7.22 | 16.85 | 13.11 | 40.11 | 20.18 | 2.16004121 | 2.94E-10 | 1.16E-08 | Up in group#2 |
| F13a1 | 7.67 | 7.79 | 12.97 | 11.6 | 28.8 | 26.76 | 68.81 | 46.84 | 2.148529706 | 1.90E-18 | 3.78E-16 | Up in group#2 |
| Mt2 | 14.11 | 9.49 | 10.43 | 8.69 | 26.5 | 31.01 | 70.15 | 52.77 | 2.137089854 | 1.14E-15 | 1.28E-13 | Up in group#2 |
| Ms4a4a | 0.9 | 1.09 | 2.33 | 1.85 | 3.81 | 4.77 | 9.69 | 7.66 | 2.123604919 | 2.91E-11 | 1.46E-09 | Up in group#2 |
| Dpep1 | 3.57 | 4.52 | 4.07 | 4.75 | 9.2 | 13.79 | 24.81 | 23.23 | 2.122232008 | 3.82E-19 | 9.08E-17 | Up in group#2 |
| Map3k6 | 6.3 | 4.59 | 6.09 | 5.95 | 15.65 | 30.16 | 25.58 | 22.61 | 2.089975719 | 1.03E-37 | 1.72E-34 | Up in group#2 |
| Inmt | 0.29 | 0.17 | 0.22 | 0.12 | 0.4 | 0.6 | 1.29 | 0.97 | 2.054935298 | 5.09E-05 | 0.000485967 | Up in group#2 |
| Ccl8 | 3.2 | 3.46 | 6.6 | 3.63 | 7.12 | 16.66 | 29.37 | 13.72 | 2.047959084 | 1.17E-08 | 3.10E-07 | Up in group#2 |
| Cd163 | 2.82 | 2.55 | 5.72 | 3.43 | 9.57 | 9.9 | 25.39 | 12.99 | 2.047784268 | 3.33E-13 | 2.39E-11 | Up in group#2 |
| Has2 | 0.32 | 0.3 | 0.12 | 0.27 | 0.6 | 0.67 | 1.26 | 1.42 | 2.014622079 | 5.65E-10 | 2.11E-08 | Up in group#2 |
| Prss35 | 0.06 | 0.06 | 0.16 | 0.17 | 0.37 | 0.25 | 0.83 | 0.31 | 1.999400757 | 3.43E-05 | 0.000346132 | Up in group#2 |
| Kcnd3 | 0.05 | 0.1 | 0.09 | 0.04 | 0.22 | 0.26 | 0.36 | 0.27 | 1.998277755 | 3.00E-10 | 1.18E-08 | Up in group#2 |
| Clec10a | 2.75 | 1.84 | 5.49 | 2.66 | 8.44 | 6.08 | 21.34 | 12.67 | 1.982222224 | 6.76E-09 | 1.91E-07 | Up in group#2 |
| Tll2 | 0.05 | 0.1 | 0.13 | 0.16 | 0.3 | 0.34 | 0.57 | 0.45 | 1.979081045 | 1.07E-07 | 2.23E-06 | Up in group#2 |
| 2010300C02F | 0.28 | 0.08 | 0.14 | 0.09 | 0.53 | 0.63 | 0.69 | 0.42 | 1.978319253 | 1.92E-09 | 6.13E-08 | Up in group#2 |
| 5031426D15I | 0.14 | 0.11 | 0.15 | 0.07 | 0.23 | 0.34 | 0.84 | 0.39 | 1.975771384 | 3.58E-07 | 6.46E-06 | Up in group#2 |
| Wnt4 | 0.09 | 0.13 | 0.15 | 0.18 | 0.49 | 0.58 | 0.57 | 0.44 | 1.969852151 | 9.60E-12 | 5.39E-10 | Up in group#2 |
| Fbn2 | 0.11 | 0.11 | 0.06 | 0.08 | 0.22 | 0.29 | 0.37 | 0.5 | 1.966436692 | 2.30E-11 | 1.18E-09 | Up in group#2 |
| Lyve1 | 5.15 | 4.79 | 10.63 | 6.2 | 19.55 | 18.69 | 39.07 | 23.22 | 1.960648848 | 3.47E-16 | 4.24E-14 | Up in group#2 |
| Apod | 2.39 | 3.41 | 3.03 | 2.54 | 8.16 | 8.7 | 12.05 | 13.56 | 1.946298001 | 3.02E-28 | 2.37E-25 | Up in group#2 |
| Mmp12 | 0.24 | 0.39 | 0.66 | 0.19 | 1.51 | 0.6 | 1.93 | 1.5 | 1.944020187 | 5.68E-08 | 1.26E-06 | Up in group#2 |
| Ereg | 0.05 | 0.05 | 0.16 | 0.08 | 0.25 | 0.25 | 0.3 | 0.45 | 1.943968895 | 1.00E-06 | 1.59E-05 | Up in group#2 |
| Susd4 | 0.36 | 0.49 | 0.3 | 0.45 | 1.2 | 1.36 | 1.8 | 1.47 | 1.936538673 | 1.85E-17 | 2.96E-15 | Up in group#2 |
| Clec4n | 0.73 | 0.77 | 1.18 | 0.91 | 2.25 | 2.18 | 4.48 | 4.07 | 1.924119416 | 1.42E-10 | 6.00E-09 | Up in group#2 |
| Clca3a1 | 0.57 | 0.83 | 0.99 | 0.72 | 2.76 | 2.82 | 3.07 | 2.76 | 1.917170878 | 2.31E-33 | 3.08E-30 | Up in group#2 |
| Penk | 0.61 | 0.84 | 0.37 | 0.99 | 1.47 | 1.89 | 3.84 | 3.16 | 1.914632502 | 3.72E-08 | 8.60E-07 | Up in group#2 |
| Lyz1 | 1.25 | 0.69 | 2.85 | 1.62 | 3.92 | 3.76 | 8.15 | 7.32 | 1.904778299 | 1.55E-08 | 3.98E-07 | Up in group#2 |
| Ccl9 | 1.73 | 1.97 | 2.59 | 2.14 | 6.07 | 3.97 | 12.33 | 8.06 | 1.897669144 | 4.81E-14 | 4.00E-12 | Up in group#2 |
| Ahsg | 2.76 | 2.69 | 13.09 | 4.61 | 24.02 | 17.13 | 25.41 | 14.84 | 1.875393321 | 3.34E-07 | 6.05E-06 | Up in group#2 |
| Mrgprb2 | 0.03 | 0.1 | 0.12 | 0.24 | 0.35 | 0.11 | 0.83 | 0.43 | 1.871476307 | 0.001171506 | 0.006828932 | Up in group#2 |
| Trnp1 | 0.1 | 0.16 | 0.44 | 0.3 | 0.72 | 0.97 | 0.98 | 0.87 | 1.862287607 | 2.75E-07 | 5.11E-06 | Up in group#2 |
| Ccr1 | 0.74 | 0.47 | 0.88 | 0.84 | 1.81 | 1.48 | 4.31 | 2.7 | 1.855901423 | 4.12E-10 | 1.57E-08 | Up in group#2 |
| Per2 | 2.24 | 2.37 | 2.44 | 1.87 | 3.27 | 11.5 | 10.07 | 6.09 | 1.848038245 | 1.21E-11 | 6.60E-10 | Up in group#2 |
| C6 | 0.18 | 0.02 | 0.12 | 0.07 | 0.16 | 0.22 | 0.71 | 0.29 | 1.8428459 | 0.000775243 | 0.004865179 | Up in group#2 |
| Tnfaip6 | 5.51 | 5.5 | 4.41 | 3.44 | 7.72 | 13.24 | 23.81 | 19.9 | 1.826431734 | 5.88E-12 | 3.40E-10 | Up in group#2 |
| Adgrg2 | 0.1 | 0.08 | 0.1 | 0.09 | 0.2 | 0.27 | 0.42 | 0.37 | 1.813321351 | 1.70E-07 | 3.38E-06 | Up in group#2 |
| Gpr160 | 2.88 | 4.16 | 4.73 | 5.69 | 11.74 | 16.74 | 17.59 | 11.91 | 1.794894296 | 5.45E-23 | 2.34E-20 | Up in group#2 |
| Mme | 0.28 | 0.27 | 0.23 | 0.19 | 0.59 | 0.67 | 0.95 | 0.95 | 1.780547729 | 1.65E-14 | 1.55E-12 | Up in group#2 |
| Sdk2 | 0.16 | 0.17 | 0.19 | 0.12 | 0.33 | 0.7 | 0.58 | 0.5 | 1.77505554 | 6.27E-11 | 2.88E-09 | Up in group#2 |
| Dpep2 | 0.2 | 0.49 | 0.41 | 0.53 | 1.02 | 1 | 1.52 | 1.3 | 1.753954437 | 2.79E-08 | 6.66E-07 | Up in group#2 |
| Mmrn1 | 1.81 | 2.42 | 2.35 | 1.59 | 9.09 | 4.47 | 7.22 | 5.55 | 1.727160769 | 2.10E-20 | 6.09E-18 | Up in group#2 |
| Fxyd2 | 1.89 | 2.01 | 2.37 | 1.78 | 5.26 | 3.42 | 10.3 | 6.78 | 1.718248637 | 3.77E-07 | 6.78E-06 | Up in group#2 |

| | | | | | | | | | | | | |
|---|---|---|---|---|---|---|---|---|---|---|---|---|
| Wisp2 | 0.48 | 0.64 | 0.69 | 0.44 | 1.36 | 1.26 | 2.24 | 2.31 | 1.706344698 | 1.85E-09 | 5.96E-08 | Up in group#2 |
| E030003E18F | 0.27 | 0.36 | 0.46 | 0.28 | 0.86 | 1.07 | 1.35 | 1.09 | 1.705947601 | 3.36E-06 | 4.63E-05 | Up in group#2 |
| Areg | 0.23 | 0.27 | 0.54 | 0.19 | 0.79 | 0.86 | 1.27 | 1.04 | 1.703449181 | 8.17E-06 | 1.00E-04 | Up in group#2 |
| Hsd11b1 | 1.81 | 3.47 | 4.21 | 2.75 | 7.75 | 6.75 | 11.72 | 12.2 | 1.700203932 | 5.27E-13 | 3.58E-11 | Up in group#2 |
| Ccl2 | 15.22 | 10.89 | 15.26 | 11.77 | 26.79 | 31.32 | 61.15 | 46.71 | 1.698237587 | 3.23E-15 | 3.40E-13 | Up in group#2 |
| Folr2 | 5.19 | 5.5 | 9.52 | 8.55 | 16.85 | 14.07 | 36.54 | 22.38 | 1.697747517 | 2.89E-11 | 1.46E-09 | Up in group#2 |
| Ctsg | 0.12 | 0.23 | 0.17 | 0.31 | 0.53 | 0.41 | 1.28 | 0.42 | 1.69763657 | 0.001712445 | 0.009273945 | Up in group#2 |
| Apoc2 | 0.63 | 0.36 | 0.51 | 0.56 | 1.35 | 1.01 | 1.51 | 2.12 | 1.688078419 | 5.69E-05 | 0.000530924 | Up in group#2 |
| Gfpt2 | 5.24 | 4.64 | 4.31 | 3.48 | 8.06 | 10.26 | 19.62 | 17.18 | 1.687802522 | 4.49E-14 | 3.78E-12 | Up in group#2 |
| Mhrt | 4.09 | 5.51 | 6.83 | 8.54 | 12.74 | 12.39 | 36.55 | 14.97 | 1.676370317 | 6.56E-08 | 1.43E-06 | Up in group#2 |
| Adamts19 | 0.05 | 0.01 | 0.15 | 0.04 | 0.09 | 0.13 | 0.3 | 0.26 | 1.664614115 | 0.002174263 | 0.011244526 | Up in group#2 |
| Kif26b | 0.06 | 0.04 | 0.11 | 0.03 | 0.15 | 0.22 | 0.25 | 0.12 | 1.663531801 | 1.57E-05 | 0.000178701 | Up in group#2 |
| Tph1 | 0.04 | 0.22 | 0.06 | 0.23 | 0.19 | 0.2 | 0.76 | 0.17 | 1.660898562 | 0.000331237 | 0.002392128 | Up in group#2 |
| Wfdc17 | 6.85 | 5.65 | 9.74 | 9.28 | 14.32 | 15.41 | 39.83 | 24.84 | 1.649748834 | 3.19E-08 | 7.55E-07 | Up in group#2 |
| Bcat1 | 0.04 | 0.05 | 0.07 | 0.04 | 0.06 | 0.16 | 0.22 | 0.18 | 1.640573997 | 4.82E-05 | 0.000462936 | Up in group#2 |
| Mlph | 0.32 | 0.2 | 0.38 | 0.31 | 0.78 | 0.92 | 1.08 | 0.88 | 1.629536756 | 8.58E-14 | 6.80E-12 | Up in group#2 |
| Slc47a2 | 0.07 | 0.15 | 0.08 | 0.14 | 0.24 | 0.51 | 0.31 | 0.23 | 1.619233902 | 0.0003684 | 0.002617918 | Up in group#2 |
| Olr1 | 0.16 | 0.23 | 0.23 | 0.18 | 0.55 | 0.59 | 0.59 | 0.63 | 1.617303078 | 7.24E-10 | 2.62E-08 | Up in group#2 |
| Tpbg | 0.23 | 0.19 | 0.08 | 0.12 | 0.19 | 0.43 | 0.68 | 0.53 | 1.616080366 | 7.94E-05 | 0.000707353 | Up in group#2 |
| Clec1b | 0.94 | 0.19 | 0.84 | 0.38 | 2.62 | 1.54 | 1.3 | 1.56 | 1.604540571 | 1.71E-05 | 0.000192929 | Up in group#2 |
| Ms4a8a | 0.38 | 0.36 | 0.46 | 0.61 | 0.81 | 0.66 | 2.47 | 1.43 | 1.603619747 | 0.000267181 | 0.001997781 | Up in group#2 |
| Phox2a | 0.24 | 0.33 | 0.13 | 0.14 | 0.24 | 0.65 | 0.92 | 0.66 | 1.599901629 | 0.000477103 | 0.003246591 | Up in group#2 |
| Thbs1 | 23.78 | 25.01 | 19.94 | 21.26 | 44.01 | 50.83 | 88.4 | 80.78 | 1.599408973 | 1.26E-19 | 3.22E-17 | Up in group#2 |
| Ccl24 | 0.61 | 0.61 | 0.7 | 1.03 | 1.95 | 0.79 | 3.41 | 2.35 | 1.580008581 | 2.20E-05 | 0.000239596 | Up in group#2 |
| Ms4a6d | 2.03 | 1.6 | 2.48 | 1.98 | 3.85 | 3.46 | 9.01 | 7.06 | 1.579928628 | 1.04E-08 | 2.80E-07 | Up in group#2 |
| C4b | 2.07 | 2.08 | 5.38 | 2.76 | 6.31 | 7.17 | 12.53 | 9.39 | 1.579441764 | 1.07E-09 | 3.69E-08 | Up in group#2 |
| Tmeff2 | 0.03 | 0.07 | 0.09 | 0.13 | 0.23 | 0.34 | 0.24 | 0.11 | 1.578806542 | 0.000752733 | 0.00474628 | Up in group#2 |
| Gm4956 | 0.25 | 0.34 | 0.43 | 0.42 | 0.55 | 1.31 | 1.27 | 1.02 | 1.577226749 | 5.71E-05 | 0.000532732 | Up in group#2 |
| Fam107a | 8.28 | 7.86 | 9.31 | 9.03 | 16.11 | 33.82 | 29.23 | 20.34 | 1.575428489 | 1.30E-18 | 2.74E-16 | Up in group#2 |
| Myoc | 0.89 | 0.98 | 0.97 | 1.05 | 1.99 | 1.94 | 4.23 | 2.99 | 1.57070762 | 4.08E-10 | 1.56E-08 | Up in group#2 |
| Rassf4 | 6.48 | 6.58 | 7.15 | 8.34 | 15.93 | 21.16 | 24.59 | 20.14 | 1.570693724 | 1.61E-40 | 3.07E-37 | Up in group#2 |
| Smarca1 | 0.65 | 0.49 | 0.65 | 0.74 | 1.52 | 1.69 | 2.04 | 1.93 | 1.562583828 | 4.91E-19 | 1.15E-16 | Up in group#2 |
| Timp1 | 0.97 | 0.73 | 1.44 | 0.64 | 1.96 | 2.27 | 3.72 | 3.01 | 1.562071054 | 9.65E-07 | 1.54E-05 | Up in group#2 |
| Tcf23 | 2.38 | 2.88 | 4.42 | 4.95 | 7.31 | 12.82 | 9.93 | 11.34 | 1.558683087 | 7.96E-15 | 7.91E-13 | Up in group#2 |
| Il1rl1 | 0.46 | 0.44 | 0.63 | 0.56 | 1.18 | 0.84 | 2.5 | 1.54 | 1.558232272 | 1.75E-08 | 4.44E-07 | Up in group#2 |
| Serpinb10 | 0.08 | 0.07 | 0.15 | 0.08 | 0.32 | 0.21 | 0.22 | 0.32 | 1.538872065 | 8.37E-05 | 0.000739019 | Up in group#2 |
| Hapln1 | 0.28 | 0.28 | 0.54 | 0.32 | 1.42 | 0.68 | 0.98 | 0.96 | 1.536403286 | 9.79E-10 | 3.41E-08 | Up in group#2 |
| Il33 | 1.12 | 1.08 | 1.09 | 0.59 | 2.86 | 2.44 | 3.73 | 1.92 | 1.536157903 | 4.30E-11 | 2.06E-09 | Up in group#2 |
| Scin | 0.09 | 0.05 | 0.11 | 0.08 | 0.14 | 0.13 | 0.27 | 0.39 | 1.530257624 | 0.001658813 | 0.009046032 | Up in group#2 |
| Srd5a2 | 0.73 | 1.46 | 0.83 | 1.2 | 1.59 | 2.96 | 2.7 | 4.49 | 1.528556999 | 1.04E-05 | 0.000124237 | Up in group#2 |
| Siglech | 0.07 | 0.14 | 0.22 | 0.26 | 0.43 | 0.24 | 0.69 | 0.56 | 1.524819838 | 0.000322415 | 0.002343669 | Up in group#2 |
| Ugt1a7c | 0.54 | 0.42 | 0.59 | 0.72 | 1.15 | 1.28 | 2.13 | 1.68 | 1.510706152 | 1.05E-10 | 4.63E-09 | Up in group#2 |

| | | | | | | | | | | | | |
|---|---|---|---|---|---|---|---|---|---|---|---|---|
| Msr1 | 1.31 | 1.08 | 1.76 | 1.42 | 3.42 | 2.51 | 6.03 | 3.62 | 1.508408458 | 1.14E-10 | 4.96E-09 | Up in group#2 |
| Mmp16 | 0.11 | 0.08 | 0.09 | 0.09 | 0.21 | 0.18 | 0.39 | 0.27 | 1.507676353 | 3.22E-05 | 0.000328783 | Up in group#2 |
| Cd209f | 0.31 | 0.5 | 0.9 | 0.7 | 0.83 | 0.82 | 3.76 | 1.25 | 1.506995885 | 0.001896035 | 0.010044418 | Up in group#2 |
| Inhba | 0.93 | 0.77 | 1.48 | 0.54 | 2.58 | 1.83 | 2.92 | 2.9 | 1.50124063 | 3.41E-11 | 1.67E-09 | Up in group#2 |
| Ccl3 | 6.45 | 3.01 | 4.36 | 6.33 | 11.79 | 8.72 | 21.97 | 12.22 | 1.496919343 | 3.72E-08 | 8.60E-07 | Up in group#2 |
| Gpr22 | 0.27 | 0.1 | 0.12 | 0.18 | 0.4 | 0.44 | 0.43 | 0.59 | 1.494104242 | 5.42E-07 | 9.21E-06 | Up in group#2 |
| Rasgef1c | 0.3 | 0.31 | 0.18 | 0.14 | 0.42 | 0.55 | 0.68 | 0.81 | 1.492333328 | 2.40E-05 | 0.000257804 | Up in group#2 |
| Lgals3 | 2.73 | 2.72 | 4.54 | 3.1 | 7.75 | 5.56 | 11.45 | 10.71 | 1.490918356 | 2.65E-11 | 1.35E-09 | Up in group#2 |
| Hp | 0.42 | 0.44 | 0.67 | 0.64 | 1.16 | 1.3 | 1.52 | 1.94 | 1.485358142 | 5.66E-07 | 9.56E-06 | Up in group#2 |
| Mrc1 | 6.24 | 5.71 | 9.16 | 7.6 | 14.58 | 11.62 | 31.21 | 20.17 | 1.484009185 | 3.33E-11 | 1.65E-09 | Up in group#2 |
| Gm44505 | 0.28 | 0.26 | 0.22 | 0.14 | 0.58 | 0.88 | 0.44 | 0.53 | 1.48339025 | 2.67E-06 | 3.79E-05 | Up in group#2 |
| Zfhx4 | 0.03 | 0.03 | 0.03 | 0.03 | 0.07 | 0.04 | 0.12 | 0.1 | 1.482027272 | 9.01E-05 | 0.000791727 | Up in group#2 |
| Gabra3 | 0.45 | 0.34 | 0.36 | 0.36 | 0.73 | 0.98 | 1.39 | 0.95 | 1.474694904 | 6.72E-10 | 2.45E-08 | Up in group#2 |
| Vit | 0.39 | 0.33 | 0.89 | 0.55 | 1.24 | 1.12 | 1.81 | 1.58 | 1.472639225 | 6.34E-08 | 1.39E-06 | Up in group#2 |
| Rgs17 | 0.14 | 0.09 | 0.06 | 0.07 | 0.17 | 0.2 | 0.31 | 0.31 | 1.471094831 | 5.39E-06 | 6.98E-05 | Up in group#2 |
| Reln | 0.47 | 0.63 | 0.47 | 0.32 | 1.46 | 1.03 | 1.38 | 1.26 | 1.46908189 | 1.42E-17 | 2.37E-15 | Up in group#2 |
| F630028O10I | 0.11 | 0.07 | 0.19 | 0.11 | 0.17 | 0.11 | 0.59 | 0.46 | 1.46696468 | 0.004685385 | 0.020847067 | Up in group#2 |
| Alox5 | 1.02 | 0.63 | 1.18 | 0.79 | 2.01 | 1.4 | 3.99 | 2.24 | 1.460241382 | 1.83E-07 | 3.57E-06 | Up in group#2 |
| Egfr | 1.06 | 1.15 | 0.98 | 1 | 1.98 | 2.18 | 3.8 | 3.57 | 1.459505981 | 1.61E-12 | 1.03E-10 | Up in group#2 |
| Ccbe1 | 0.09 | 0.08 | 0.13 | 0.11 | 0.23 | 0.23 | 0.38 | 0.27 | 1.45629817 | 1.94E-06 | 2.88E-05 | Up in group#2 |
| Cdh19 | 1.17 | 1.19 | 1.17 | 1.33 | 2.26 | 2.34 | 4.26 | 4.04 | 1.449465495 | 6.95E-13 | 4.56E-11 | Up in group#2 |
| Slc10a6 | 4.68 | 4.89 | 5.56 | 5.91 | 10.44 | 12.34 | 19.94 | 12.62 | 1.446124065 | 1.09E-17 | 1.83E-15 | Up in group#2 |
| Eya2 | 0.37 | 0.26 | 0.15 | 0.26 | 0.46 | 0.53 | 0.86 | 0.85 | 1.442481028 | 2.07E-05 | 0.000227316 | Up in group#2 |
| Lyz2 | 102.86 | 89.34 | 154.2 | 115.01 | 244.63 | 200.49 | 418.29 | 344.31 | 1.441656182 | 3.37E-14 | 2.98E-12 | Up in group#2 |
| Hpgd | 2.32 | 2.84 | 4.7 | 3.47 | 8.03 | 6.35 | 12.08 | 8.51 | 1.44048977 | 5.93E-12 | 3.42E-10 | Up in group#2 |
| Cyp11a1 | 0.36 | 0.12 | 0.23 | 0.37 | 0.67 | 0.61 | 0.79 | 0.7 | 1.437811738 | 2.96E-06 | 4.14E-05 | Up in group#2 |
| Slc45a3 | 1.2 | 0.98 | 2.17 | 0.92 | 3.27 | 2.92 | 3.9 | 3.67 | 1.434523104 | 4.23E-11 | 2.03E-09 | Up in group#2 |
| C3 | 5.15 | 3.94 | 8.81 | 4.64 | 8.55 | 10.19 | 23.29 | 16.71 | 1.434239055 | 9.34E-08 | 1.98E-06 | Up in group#2 |
| Ccl4 | 4.44 | 1.77 | 3.22 | 2.67 | 5.75 | 5.35 | 10.2 | 10.17 | 1.433620937 | 8.04E-07 | 1.30E-05 | Up in group#2 |
| Anxa8 | 0.27 | 0.36 | 0.51 | 0.22 | 0.62 | 0.67 | 1.49 | 0.73 | 1.431054708 | 0.00014638 | 0.001201076 | Up in group#2 |
| Omd | 0.4 | 0.5 | 0.33 | 0.27 | 0.6 | 1.22 | 1.07 | 0.99 | 1.425147352 | 8.30E-06 | 0.000101404 | Up in group#2 |
| Fam167a | 0.2 | 0.26 | 0.25 | 0.29 | 0.66 | 0.62 | 0.69 | 0.65 | 1.42430736 | 9.06E-11 | 4.04E-09 | Up in group#2 |
| Adamtsl2 | 1.94 | 1.91 | 1.48 | 1.05 | 3.06 | 2.93 | 5.91 | 4.72 | 1.422786305 | 9.02E-10 | 3.18E-08 | Up in group#2 |
| Stra6 | 0.2 | 0.2 | 0.32 | 0.23 | 0.54 | 0.61 | 0.82 | 0.51 | 1.422729467 | 1.93E-06 | 2.88E-05 | Up in group#2 |
| Srpx2 | 0.85 | 1.31 | 0.79 | 0.97 | 2 | 2.08 | 3.52 | 2.49 | 1.415026619 | 9.06E-11 | 4.04E-09 | Up in group#2 |
| Lif | 0.18 | 0.13 | 0.26 | 0.14 | 0.5 | 0.38 | 0.59 | 0.4 | 1.414286431 | 1.16E-06 | 1.81E-05 | Up in group#2 |
| Tnfaip8l3 | 0.09 | 0.16 | 0.34 | 0.1 | 0.54 | 0.35 | 0.49 | 0.44 | 1.414096295 | 0.000360385 | 0.002571945 | Up in group#2 |
| Agap2 | 1.48 | 1.37 | 1.32 | 1.75 | 2.6 | 3.61 | 4.83 | 4.18 | 1.411486595 | 4.90E-17 | 6.94E-15 | Up in group#2 |
| Tbx1 | 0.33 | 0.35 | 0.55 | 0.19 | 0.99 | 0.81 | 0.97 | 0.88 | 1.408786872 | 4.46E-06 | 5.95E-05 | Up in group#2 |
| Chrd | 0.16 | 0.18 | 0.34 | 0.27 | 0.4 | 0.53 | 0.8 | 0.74 | 1.407401519 | 1.04E-05 | 0.000124237 | Up in group#2 |
| Fbp2 | 0.3 | 0.38 | 0.75 | 0.81 | 1.3 | 1.81 | 0.94 | 1.66 | 1.404358638 | 3.55E-05 | 0.000356365 | Up in group#2 |
| Gdf6 | 0.17 | 0.06 | 0.05 | 0.03 | 0.09 | 0.15 | 0.17 | 0.39 | 1.402930271 | 0.011087943 | 0.041291425 | Up in group#2 |

| | | | | | | | | | | | | |
|---|---|---|---|---|---|---|---|---|---|---|---|---|
| Ace2 | 1.33 | 1.04 | 0.99 | 1.45 | 2.36 | 2.38 | 4 | 3.56 | 1.400354363 | 1.94E-12 | 1.23E-10 | Up in group#2 |
| Myc | 9.05 | 7.86 | 12.69 | 7.64 | 19.7 | 21.8 | 29.33 | 23.76 | 1.394948017 | 1.11E-20 | 3.42E-18 | Up in group#2 |
| P2ry12 | 0.75 | 0.61 | 1.46 | 1.37 | 2.99 | 1.77 | 3.65 | 2.33 | 1.394131867 | 1.96E-07 | 3.79E-06 | Up in group#2 |
| Aoah | 0.39 | 0.37 | 0.69 | 0.52 | 0.99 | 0.94 | 1.84 | 1.45 | 1.392899015 | 2.71E-07 | 5.06E-06 | Up in group#2 |
| Ildr2 | 0.12 | 0.13 | 0.19 | 0.11 | 0.34 | 0.27 | 0.46 | 0.33 | 1.392185837 | 7.81E-08 | 1.68E-06 | Up in group#2 |
| Kcne4 | 3.2 | 2.69 | 3.04 | 2.5 | 5.53 | 6.2 | 8.63 | 8.51 | 1.385778819 | 5.18E-18 | 9.71E-16 | Up in group#2 |
| Cbr2 | 8.12 | 8.29 | 13.95 | 8.93 | 17.66 | 14.53 | 42.88 | 23.73 | 1.38326869 | 2.41E-07 | 4.55E-06 | Up in group#2 |
| Clec4d | 1.3 | 0.89 | 1.72 | 1.26 | 2.17 | 2 | 5.02 | 3.83 | 1.383121953 | 4.23E-06 | 5.68E-05 | Up in group#2 |
| Tlr5 | 0.16 | 0.06 | 0.17 | 0.16 | 0.33 | 0.15 | 0.52 | 0.4 | 1.381276747 | 0.000865475 | 0.005343317 | Up in group#2 |
| Sele | 5.86 | 4.77 | 6.31 | 4.21 | 11.31 | 12.19 | 12.11 | 17.66 | 1.381058968 | 6.70E-20 | 1.78E-17 | Up in group#2 |
| P2ry13 | 0.23 | 0.43 | 0.67 | 0.55 | 1.29 | 0.5 | 1.81 | 1.15 | 1.37575706 | 0.000107907 | 0.000925303 | Up in group#2 |
| Ltbp2 | 0.3 | 0.3 | 0.28 | 0.17 | 0.76 | 0.54 | 0.81 | 0.54 | 1.375402034 | 1.06E-09 | 3.67E-08 | Up in group#2 |
| Nupr1 | 4.12 | 4.67 | 6.11 | 4.5 | 12.09 | 10.4 | 16.23 | 9.89 | 1.373159919 | 5.79E-11 | 2.68E-09 | Up in group#2 |
| C5ar1 | 3.14 | 2.32 | 3.13 | 2.59 | 5.54 | 3.74 | 11.07 | 7.63 | 1.373104989 | 1.60E-08 | 4.10E-07 | Up in group#2 |
| Rbm47 | 0.15 | 0.07 | 0.1 | 0.05 | 0.2 | 0.14 | 0.31 | 0.26 | 1.368661761 | 0.000326966 | 0.0023677 | Up in group#2 |
| Frzb | 1.32 | 1.66 | 1.42 | 1.14 | 3.21 | 2.71 | 4.08 | 3.85 | 1.366224315 | 1.66E-15 | 1.81E-13 | Up in group#2 |
| 6030408B16I | 0.11 | 0.21 | 0.15 | 0.25 | 0.24 | 0.23 | 0.68 | 0.65 | 1.362920201 | 0.003270902 | 0.01566293 | Up in group#2 |
| Dnm1 | 0.79 | 0.93 | 1.14 | 0.76 | 1.73 | 1.37 | 3.37 | 2.63 | 1.361963161 | 3.89E-08 | 8.92E-07 | Up in group#2 |
| Kcnk13 | 0.17 | 0.32 | 0.24 | 0.19 | 0.42 | 0.29 | 1.21 | 0.37 | 1.357873396 | 0.001610562 | 0.008808154 | Up in group#2 |
| Akr1b8 | 1.05 | 1.3 | 1.49 | 0.7 | 2.52 | 2.34 | 3.63 | 2.8 | 1.354134231 | 3.56E-08 | 8.31E-07 | Up in group#2 |
| Rab27b | 0.19 | 0.16 | 0.25 | 0.23 | 0.6 | 0.39 | 0.68 | 0.38 | 1.353610616 | 4.58E-08 | 1.03E-06 | Up in group#2 |
| Gsn | 245.1 | 245.76 | 272.98 | 200.96 | 403.08 | 440.75 | 840.75 | 696.31 | 1.352309616 | 5.42E-13 | 3.65E-11 | Up in group#2 |
| S100a4 | 9.81 | 8.25 | 13.56 | 9.35 | 17.95 | 21.26 | 37.01 | 23.66 | 1.346881441 | 4.68E-09 | 1.38E-07 | Up in group#2 |
| Cldn15 | 10.19 | 10.15 | 12.06 | 10.62 | 27.51 | 24.73 | 28.24 | 24.68 | 1.337568039 | 1.56E-50 | 1.04E-46 | Up in group#2 |
| Thbs4 | 0.37 | 0.69 | 0.23 | 0.38 | 1.03 | 1.04 | 1.42 | 0.59 | 1.334028031 | 2.82E-05 | 0.000295155 | Up in group#2 |
| Cadm2 | 0.07 | 0.09 | 0.05 | 0.05 | 0.12 | 0.13 | 0.21 | 0.21 | 1.332427697 | 2.88E-05 | 0.000300335 | Up in group#2 |
| Ccl6 | 9.87 | 9.68 | 15.75 | 17.65 | 24.56 | 19.24 | 50.72 | 33.84 | 1.331864091 | 4.79E-08 | 1.07E-06 | Up in group#2 |
| Meg3 | 3.22 | 2.58 | 2.77 | 3 | 5.94 | 5.12 | 9.12 | 6.03 | 1.330836754 | 8.22E-18 | 1.44E-15 | Up in group#2 |
| Klf15 | 1.3 | 1.3 | 1.32 | 1.03 | 1.83 | 3.58 | 3.67 | 2.74 | 1.329074332 | 7.21E-10 | 2.62E-08 | Up in group#2 |
| Lrg1 | 4.59 | 3.77 | 4.17 | 2.86 | 9.5 | 8.68 | 10.16 | 9.04 | 1.324967151 | 8.82E-19 | 1.92E-16 | Up in group#2 |
| Htr2b | 0.08 | 0.27 | 0.17 | 0.16 | 0.58 | 0.2 | 0.48 | 0.41 | 1.318807064 | 0.00247683 | 0.012508134 | Up in group#2 |
| Il1rn | 1.42 | 0.99 | 1.68 | 1.36 | 2.79 | 2.31 | 4.28 | 3.73 | 1.312189283 | 4.16E-10 | 1.58E-08 | Up in group#2 |
| Flrt3 | 0.46 | 0.34 | 0.61 | 0.34 | 0.98 | 0.82 | 1.33 | 1.06 | 1.309333197 | 9.88E-09 | 2.68E-07 | Up in group#2 |
| Cyp1b1 | 1.17 | 1.08 | 2.44 | 0.98 | 2.48 | 2.71 | 4.43 | 3.96 | 1.30710099 | 1.82E-07 | 3.56E-06 | Up in group#2 |
| Clec12b | 3.78 | 4.3 | 4.39 | 5.36 | 8.07 | 11.79 | 11.9 | 10.69 | 1.306611409 | 4.28E-17 | 6.13E-15 | Up in group#2 |
| Mfap5 | 7.06 | 5.51 | 7.61 | 4.87 | 11.26 | 13.36 | 21.65 | 13.46 | 1.305279132 | 1.21E-11 | 6.60E-10 | Up in group#2 |
| Dchs2 | 0.04 | 0.02 | 0.04 | 0.03 | 0.06 | 0.06 | 0.09 | 0.1 | 1.303059949 | 0.000710779 | 0.004507356 | Up in group#2 |
| Rnase4 | 17.78 | 19.76 | 24.47 | 16.74 | 39.62 | 37.93 | 62.01 | 47.98 | 1.300699618 | 5.45E-19 | 1.25E-16 | Up in group#2 |
| Ackr4 | 1.02 | 0.87 | 1.19 | 0.83 | 2.29 | 1.48 | 3.37 | 2.19 | 1.296461038 | 1.04E-07 | 2.17E-06 | Up in group#2 |
| Bdnf | 0.16 | 0.19 | 0.19 | 0.12 | 0.23 | 0.28 | 0.67 | 0.4 | 1.291978059 | 0.00047261 | 0.003222606 | Up in group#2 |
| Trf | 13.49 | 11.12 | 20.4 | 14.14 | 27.35 | 23.58 | 50.74 | 37.95 | 1.289719424 | 1.70E-10 | 7.08E-09 | Up in group#2 |
| Adamts5 | 7.72 | 6.67 | 7.11 | 7.28 | 15.83 | 14.85 | 19.41 | 17.87 | 1.287361348 | 9.24E-48 | 3.42E-44 | Up in group#2 |

| | | | | | | | | | | | | |
|---|---|---|---|---|---|---|---|---|---|---|---|---|
| C1qa | 29.9 | 20.83 | 38.61 | 27.28 | 47.66 | 47.85 | 107.23 | 71.32 | 1.287003753 | 5.46E-09 | 1.59E-07 | Up in group#2 |
| Wif1 | 0.12 | 0.25 | 0.36 | 0.05 | 0.51 | 0.32 | 0.66 | 0.35 | 1.278256828 | 0.003474939 | 0.016462383 | Up in group#2 |
| Ctgf | 20.33 | 26.53 | 30.48 | 28.41 | 52.89 | 53.67 | 68.25 | 72.33 | 1.275195682 | 1.51E-25 | 8.06E-23 | Up in group#2 |
| Ms4a6c | 1.95 | 1.66 | 3.36 | 2.72 | 4.35 | 4.17 | 9.28 | 7.24 | 1.274523715 | 1.93E-06 | 2.88E-05 | Up in group#2 |
| Rian | 1.95 | 1.77 | 1.52 | 1.85 | 3.46 | 3.23 | 5.03 | 4.89 | 1.274498542 | 6.30E-15 | 6.46E-13 | Up in group#2 |
| Fcgr3 | 8.64 | 6.73 | 12.57 | 9.65 | 17.65 | 15.54 | 31.5 | 22.96 | 1.272414208 | 2.91E-10 | 1.15E-08 | Up in group#2 |
| Cilp | 0.64 | 0.64 | 0.53 | 0.45 | 1.32 | 1.27 | 1.69 | 1.02 | 1.27202029 | 3.21E-10 | 1.25E-08 | Up in group#2 |
| Timp4 | 79.5 | 77.39 | 85.2 | 113.59 | 147.77 | 287.17 | 209.53 | 180.71 | 1.271158265 | 6.84E-14 | 5.52E-12 | Up in group#2 |
| Stk32b | 0.21 | 0.17 | 0.24 | 0.11 | 0.29 | 0.4 | 0.61 | 0.39 | 1.264747565 | 0.000161274 | 0.001307177 | Up in group#2 |
| Scn7a | 4.57 | 4.79 | 4.31 | 4.18 | 7.58 | 7.39 | 13.64 | 12.83 | 1.261766396 | 4.08E-14 | 3.53E-12 | Up in group#2 |
| Mt1 | 44.62 | 35.79 | 33.98 | 35.01 | 51.69 | 69.3 | 114.6 | 107.81 | 1.261662094 | 4.78E-10 | 1.80E-08 | Up in group#2 |
| Fam105a | 0.84 | 0.58 | 0.99 | 0.67 | 1.49 | 1.24 | 2.66 | 1.77 | 1.261471303 | 2.80E-07 | 5.18E-06 | Up in group#2 |
| Slc7a8 | 0.82 | 0.78 | 1.02 | 0.72 | 1.61 | 1.05 | 3.06 | 2.05 | 1.259614753 | 9.31E-07 | 1.49E-05 | Up in group#2 |
| Mafa | 2.16 | 2.28 | 2.37 | 2.8 | 4.36 | 6.62 | 5.04 | 6.08 | 1.258082571 | 1.59E-10 | 6.69E-09 | Up in group#2 |
| Agtr1a | 1.19 | 1.21 | 1.03 | 1.32 | 2.64 | 2.15 | 3.2 | 3.01 | 1.256682285 | 3.59E-12 | 2.15E-10 | Up in group#2 |
| Tnfsf14 | 0.14 | 0.22 | 0.27 | 0.09 | 0.34 | 0.27 | 0.43 | 0.66 | 1.256313628 | 0.004017161 | 0.018491716 | Up in group#2 |
| Rerg | 1.91 | 1.72 | 1.24 | 1.77 | 2.94 | 3.37 | 4.9 | 4.06 | 1.255275771 | 5.74E-11 | 2.67E-09 | Up in group#2 |
| Slc9a9 | 1.09 | 0.77 | 1.36 | 1.21 | 2.06 | 1.64 | 3.98 | 2.54 | 1.254958416 | 2.50E-07 | 4.72E-06 | Up in group#2 |
| Slc38a4 | 0.34 | 0.45 | 0.62 | 0.41 | 1.07 | 0.96 | 1.22 | 0.94 | 1.249679612 | 2.01E-09 | 6.34E-08 | Up in group#2 |
| Gna14 | 0.36 | 0.12 | 0.31 | 0.23 | 0.52 | 0.52 | 0.55 | 0.76 | 1.24817877 | 3.43E-05 | 0.000346526 | Up in group#2 |
| Gsg1l | 0.09 | 0.16 | 0.14 | 0.11 | 0.29 | 0.26 | 0.3 | 0.26 | 1.247035333 | 0.000108533 | 0.000930073 | Up in group#2 |
| Kcnj10 | 0.12 | 0.05 | 0.07 | 0.06 | 0.13 | 0.14 | 0.17 | 0.24 | 1.24288373 | 0.001210325 | 0.006983492 | Up in group#2 |
| Mst1r | 0.15 | 0.37 | 0.26 | 0.13 | 0.39 | 0.57 | 0.76 | 0.55 | 1.239398696 | 6.31E-05 | 0.000580353 | Up in group#2 |
| Timd4 | 0.89 | 0.74 | 1.77 | 1.02 | 2.07 | 1.34 | 3.78 | 2.93 | 1.237747447 | 4.01E-05 | 0.000396646 | Up in group#2 |
| Dnm3os | 0.38 | 0.35 | 0.28 | 0.32 | 0.44 | 0.61 | 1.03 | 0.94 | 1.237728209 | 4.52E-07 | 7.89E-06 | Up in group#2 |
| Fcgr2b | 5.74 | 5.43 | 7.89 | 7.57 | 10.99 | 11.49 | 21.28 | 16.9 | 1.231590279 | 1.73E-10 | 7.21E-09 | Up in group#2 |
| Colec11 | 20.37 | 19.47 | 21.55 | 21.85 | 41.13 | 38.17 | 54.24 | 54.97 | 1.230225169 | 1.06E-27 | 7.40E-25 | Up in group#2 |
| Ticam2 | 0.53 | 0.3 | 0.39 | 0.34 | 0.81 | 0.83 | 1.1 | 0.8 | 1.228793238 | 2.83E-07 | 5.21E-06 | Up in group#2 |
| Pf4 | 16.26 | 12.45 | 20.79 | 19.87 | 32.24 | 26.91 | 55.89 | 40.67 | 1.226328258 | 2.50E-09 | 7.75E-08 | Up in group#2 |
| Hs3st3a1 | 0.19 | 0.14 | 0.11 | 0.07 | 0.29 | 0.27 | 0.4 | 0.19 | 1.220528356 | 0.000892996 | 0.005484026 | Up in group#2 |
| Foxp2 | 0.29 | 0.49 | 0.35 | 0.25 | 0.83 | 0.73 | 1.01 | 0.63 | 1.219715407 | 3.97E-08 | 9.09E-07 | Up in group#2 |
| Ms4a2 | 0.11 | 0.08 | 0.11 | 0.17 | 0.19 | 0.17 | 0.46 | 0.25 | 1.219573488 | 0.00723971 | 0.029375751 | Up in group#2 |
| Slc1a3 | 0.27 | 0.22 | 0.25 | 0.15 | 0.43 | 0.38 | 0.58 | 0.62 | 1.21874865 | 1.12E-05 | 0.000131711 | Up in group#2 |
| C1qc | 31.38 | 22.09 | 35.52 | 27.31 | 51.05 | 45.58 | 100.26 | 63.98 | 1.218202018 | 5.68E-10 | 2.12E-08 | Up in group#2 |
| Sfrp2 | 0.64 | 0.76 | 1.11 | 0.63 | 1.5 | 0.99 | 2.7 | 1.9 | 1.216955832 | 3.96E-05 | 0.00039293 | Up in group#2 |
| Pdpn | 1.51 | 1.09 | 1.72 | 1.11 | 2.52 | 2.35 | 4.11 | 3.25 | 1.216048898 | 4.33E-08 | 9.83E-07 | Up in group#2 |
| Htr7 | 0.51 | 0.52 | 0.3 | 0.47 | 0.84 | 1.22 | 1.37 | 0.96 | 1.215260381 | 4.83E-07 | 8.34E-06 | Up in group#2 |
| Ptgis | 3.2 | 4.23 | 6.18 | 2.86 | 8.66 | 7.21 | 11.28 | 9.89 | 1.215242962 | 2.06E-09 | 6.48E-08 | Up in group#2 |
| Tenm3 | 0.32 | 0.33 | 0.26 | 0.21 | 0.54 | 0.41 | 0.85 | 0.71 | 1.210853054 | 6.70E-08 | 1.46E-06 | Up in group#2 |
| Gas7 | 1.68 | 1.4 | 1.71 | 1.42 | 2.88 | 2.3 | 4.8 | 3.82 | 1.208769352 | 1.50E-11 | 7.96E-10 | Up in group#2 |
| Apba2 | 0.14 | 0.21 | 0.09 | 0.26 | 0.26 | 0.35 | 0.45 | 0.48 | 1.206909085 | 0.000616962 | 0.004021579 | Up in group#2 |
| Nrcam | 0.12 | 0.17 | 0.17 | 0.1 | 0.31 | 0.26 | 0.46 | 0.24 | 1.206459941 | 1.86E-05 | 0.000208365 | Up in group#2 |

| | | | | | | | | | | | | |
|---|---|---|---|---|---|---|---|---|---|---|---|---|
| Tubb1 | 1.14 | 0.51 | 1.01 | 1.86 | 3.29 | 2.18 | 1.68 | 2.84 | 1.199268149 | 3.38E-05 | 0.000342229 | Up in group#2 |
| LOC1000389 | 0.23 | 0.29 | 0.54 | 0.16 | 0.53 | 0.56 | 0.71 | 0.94 | 1.19911719 | 0.002637198 | 0.01317313 | Up in group#2 |
| Fmn1 | 0.08 | 0.08 | 0.16 | 0.05 | 0.17 | 0.13 | 0.28 | 0.21 | 1.199023439 | 0.000482761 | 0.003272501 | Up in group#2 |
| Retnla | 20.15 | 20.76 | 45.96 | 30.23 | 43.29 | 49.82 | 104.99 | 59.14 | 1.19861939 | 7.18E-06 | 8.93E-05 | Up in group#2 |
| Htra3 | 9.38 | 7.16 | 8.16 | 6.62 | 11.54 | 13.6 | 23.38 | 21.02 | 1.195624015 | 9.42E-11 | 4.18E-09 | Up in group#2 |
| Camk2n1 | 11.08 | 11.1 | 11.49 | 12.27 | 22.99 | 22.5 | 28.94 | 26.66 | 1.186365228 | 6.40E-41 | 1.42E-37 | Up in group#2 |
| Rhou | 0.54 | 0.46 | 0.75 | 0.51 | 0.99 | 1.23 | 1.56 | 1.22 | 1.185997768 | 3.22E-08 | 7.58E-07 | Up in group#2 |
| Arnt2 | 0.06 | 0.09 | 0.19 | 0.16 | 0.26 | 0.23 | 0.34 | 0.27 | 1.184404791 | 0.00024247 | 0.001843022 | Up in group#2 |
| Hsph1 | 55.96 | 42.06 | 28.12 | 24.88 | 71.31 | 100.4 | 76.58 | 84.35 | 1.180992175 | 3.11E-09 | 9.44E-08 | Up in group#2 |
| Gpr34 | 1.43 | 1.32 | 1.42 | 1.54 | 3.27 | 1.83 | 4.92 | 2.54 | 1.179352264 | 2.84E-06 | 3.99E-05 | Up in group#2 |
| Cd33 | 1.15 | 1.08 | 1.35 | 0.97 | 1.9 | 1.82 | 3.71 | 2.53 | 1.17907103 | 2.71E-07 | 5.06E-06 | Up in group#2 |
| Il6 | 30.12 | 20.08 | 19.55 | 18.83 | 37.48 | 38.44 | 62.01 | 55.45 | 1.175533723 | 1.09E-11 | 6.04E-10 | Up in group#2 |
| Cd14 | 19.44 | 12.9 | 20.55 | 14.83 | 31.83 | 27.2 | 52.07 | 36.7 | 1.175086139 | 1.16E-11 | 6.40E-10 | Up in group#2 |
| C1qb | 27.56 | 22.42 | 36.92 | 25.06 | 44.59 | 42.29 | 93.99 | 62.26 | 1.171896625 | 1.58E-08 | 4.05E-07 | Up in group#2 |
| Mmp19 | 1.23 | 1.38 | 1.11 | 1.34 | 2.39 | 1.94 | 3.54 | 3.03 | 1.171864664 | 3.29E-10 | 1.28E-08 | Up in group#2 |
| Cp | 13.38 | 14.18 | 21.32 | 11.49 | 30.01 | 27.71 | 39.81 | 34.38 | 1.169345154 | 7.11E-15 | 7.22E-13 | Up in group#2 |
| Npl | 0.99 | 0.46 | 0.98 | 1.15 | 1.54 | 1.75 | 2.41 | 2.03 | 1.168102233 | 1.42E-05 | 0.000163068 | Up in group#2 |
| Tgfbi | 7.37 | 6.94 | 11.54 | 8.01 | 14.68 | 14.71 | 26.6 | 17.4 | 1.166413495 | 1.00E-10 | 4.43E-09 | Up in group#2 |
| Plekhb1 | 1.73 | 2.09 | 2.54 | 1.85 | 3.66 | 3.64 | 5.67 | 4.89 | 1.166141183 | 8.54E-10 | 3.04E-08 | Up in group#2 |
| Proser2 | 0.07 | 0.14 | 0.28 | 0.2 | 0.35 | 0.4 | 0.37 | 0.36 | 1.166056284 | 0.000464716 | 0.003176908 | Up in group#2 |
| Krt19 | 0.28 | 0.38 | 0.93 | 0.55 | 1.18 | 0.7 | 2.1 | 0.83 | 1.164682612 | 0.00418089 | 0.019087045 | Up in group#2 |
| Tiam2 | 0.16 | 0.16 | 0.21 | 0.13 | 0.2 | 0.26 | 0.41 | 0.31 | 1.164070989 | 0.00015765 | 0.001282215 | Up in group#2 |
| Srpx | 0.57 | 0.57 | 0.44 | 0.48 | 1.05 | 1.03 | 1.27 | 1.13 | 1.160055171 | 1.48E-07 | 2.96E-06 | Up in group#2 |
| Kif21a | 0.3 | 0.22 | 0.48 | 0.4 | 0.76 | 0.64 | 0.9 | 0.73 | 1.15741665 | 5.21E-08 | 1.16E-06 | Up in group#2 |
| Ifitm10 | 1.44 | 1.8 | 1.37 | 2.03 | 3.09 | 3.21 | 4.28 | 3.79 | 1.155226697 | 1.24E-08 | 3.27E-07 | Up in group#2 |
| Cytl1 | 19.49 | 19.16 | 46.28 | 18.12 | 56.31 | 54.69 | 60.33 | 49.55 | 1.153087555 | 2.68E-07 | 5.03E-06 | Up in group#2 |
| Bche | 0.08 | 0.12 | 0.18 | 0.11 | 0.24 | 0.28 | 0.26 | 0.26 | 1.150418437 | 5.24E-05 | 0.000495547 | Up in group#2 |
| Osm | 1.45 | 1.05 | 1.52 | 1.38 | 2.39 | 2.01 | 4.13 | 3.05 | 1.147870185 | 5.25E-07 | 8.99E-06 | Up in group#2 |
| Serpine2 | 9.28 | 8.34 | 7.97 | 7.75 | 15.94 | 14.47 | 22.14 | 18.86 | 1.146097394 | 1.06E-19 | 2.78E-17 | Up in group#2 |
| Plaur | 36.54 | 29.54 | 24.47 | 24.88 | 45.12 | 71.62 | 55.72 | 74.05 | 1.145915884 | 2.32E-13 | 1.74E-11 | Up in group#2 |
| Gas1 | 3.47 | 3.55 | 3.93 | 2.99 | 5.22 | 5.62 | 9.25 | 9.74 | 1.145592711 | 5.70E-10 | 2.12E-08 | Up in group#2 |
| Fst | 0.22 | 0.3 | 0.24 | 0.22 | 0.55 | 0.33 | 0.86 | 0.4 | 1.143900633 | 0.000840723 | 0.005229289 | Up in group#2 |
| Clec5a | 0.43 | 0.25 | 0.31 | 0.25 | 0.58 | 0.44 | 0.89 | 0.74 | 1.138546545 | 4.87E-05 | 0.000466331 | Up in group#2 |
| Abca6 | 1.68 | 1.71 | 1.59 | 1.53 | 2.34 | 2.82 | 4.75 | 4.29 | 1.134388986 | 7.27E-09 | 2.04E-07 | Up in group#2 |
| Pdgfc | 0.34 | 0.29 | 0.77 | 0.28 | 0.84 | 0.56 | 1.33 | 0.85 | 1.133470924 | 0.000483234 | 0.00327326 | Up in group#2 |
| Gpx3 | 21.26 | 20.62 | 28.16 | 20.78 | 32.19 | 37.25 | 64.51 | 57.69 | 1.128917404 | 2.81E-10 | 1.12E-08 | Up in group#2 |
| Maf | 2.16 | 2.02 | 2.45 | 2.12 | 3.95 | 2.98 | 6.8 | 4.8 | 1.128249315 | 5.93E-09 | 1.69E-07 | Up in group#2 |
| Megf10 | 0.16 | 0.23 | 0.22 | 0.09 | 0.34 | 0.22 | 0.55 | 0.4 | 1.126877384 | 0.000227092 | 0.001741038 | Up in group#2 |
| Chl1 | 0.4 | 0.38 | 0.37 | 0.31 | 0.64 | 0.52 | 0.96 | 1 | 1.126495165 | 2.09E-07 | 4.02E-06 | Up in group#2 |
| Tcf21 | 3.9 | 4.77 | 5.08 | 5.03 | 6.88 | 7.73 | 11.65 | 13.18 | 1.123025778 | 1.83E-08 | 4.58E-07 | Up in group#2 |
| Lilr4b | 4.33 | 2.49 | 4.62 | 3.71 | 6.54 | 5.67 | 10.91 | 8.46 | 1.120400406 | 8.71E-08 | 1.85E-06 | Up in group#2 |
| Wnt5a | 0.13 | 0.29 | 0.19 | 0.07 | 0.35 | 0.3 | 0.48 | 0.26 | 1.120293842 | 0.001030932 | 0.006195361 | Up in group#2 |

| | | | | | | | | | | | | |
|---|---|---|---|---|---|---|---|---|---|---|---|---|
| Col12a1 | 0.18 | 0.31 | 0.32 | 0.2 | 0.5 | 0.41 | 0.58 | 0.64 | 1.119789684 | 8.05E-08 | 1.72E-06 | Up in group#2 |
| Serpinb9b | 0.55 | 0.58 | 0.66 | 0.6 | 1.15 | 1.01 | 1.78 | 1.1 | 1.118260063 | 2.01E-05 | 0.000221645 | Up in group#2 |
| Glt28d2 | 0.77 | 0.71 | 0.75 | 1.05 | 1.4 | 1.72 | 2.14 | 1.6 | 1.116925424 | 3.42E-08 | 8.04E-07 | Up in group#2 |
| Gabra1 | 0.11 | 0.18 | 0.16 | 0.1 | 0.34 | 0.32 | 0.25 | 0.24 | 1.116374521 | 0.000410595 | 0.002876326 | Up in group#2 |
| Gria3 | 0.14 | 0.13 | 0.13 | 0.19 | 0.26 | 0.26 | 0.41 | 0.31 | 1.116155715 | 0.000112518 | 0.000960513 | Up in group#2 |
| Kcp | 0.36 | 0.29 | 0.24 | 0.2 | 0.45 | 0.45 | 0.91 | 0.5 | 1.11554904 | 9.56E-05 | 0.000832872 | Up in group#2 |
| Hs6st2 | 0.43 | 0.49 | 0.66 | 0.59 | 0.98 | 1.26 | 1.14 | 1.1 | 1.114771503 | 7.50E-10 | 2.70E-08 | Up in group#2 |
| Maff | 38.04 | 31.95 | 26.77 | 28.97 | 57.29 | 67.2 | 66.48 | 71.86 | 1.113891122 | 1.18E-24 | 5.80E-22 | Up in group#2 |
| Pla2g7 | 7.95 | 8.33 | 12.54 | 12.61 | 19.68 | 15.82 | 27.79 | 23.2 | 1.113569574 | 3.22E-11 | 1.61E-09 | Up in group#2 |
| 8430408G22I | 54.51 | 73.39 | 80.19 | 97.97 | 141.73 | 165.96 | 185.04 | 144.54 | 1.112649892 | 9.18E-17 | 1.23E-14 | Up in group#2 |
| Lpl | 236.67 | 247.17 | 265.9 | 383.91 | 438.95 | 693.07 | 654.48 | 567.28 | 1.108858864 | 2.90E-12 | 1.76E-10 | Up in group#2 |
| Edil3 | 0.19 | 0.27 | 0.23 | 0.15 | 0.46 | 0.37 | 0.46 | 0.49 | 1.108231097 | 3.54E-06 | 4.85E-05 | Up in group#2 |
| Cdc42ep4 | 37.41 | 33.05 | 32.87 | 31.1 | 55.79 | 71.92 | 80.31 | 72.91 | 1.106770789 | 1.18E-25 | 6.67E-23 | Up in group#2 |
| Lsr | 1.58 | 1.85 | 3.11 | 2.07 | 4.68 | 4.13 | 4.49 | 4.42 | 1.106351163 | 5.57E-10 | 2.09E-08 | Up in group#2 |
| Grik3 | 0.09 | 0.07 | 0.13 | 0.12 | 0.17 | 0.13 | 0.28 | 0.29 | 1.102830559 | 0.000374447 | 0.002656638 | Up in group#2 |
| Csf1r | 11.99 | 11 | 18.21 | 12.39 | 22.71 | 18.68 | 40.19 | 29.54 | 1.100954409 | 5.77E-09 | 1.65E-07 | Up in group#2 |
| Gsta3 | 1.8 | 2.59 | 2.33 | 1.61 | 3.53 | 3.39 | 5.8 | 4.76 | 1.100457046 | 4.02E-07 | 7.15E-06 | Up in group#2 |
| Serpine1 | 19.59 | 16.53 | 14.01 | 14.14 | 25.89 | 30.6 | 40.5 | 36.28 | 1.09966764 | 4.69E-16 | 5.58E-14 | Up in group#2 |
| Sulf2 | 3.43 | 2.99 | 5.35 | 3.29 | 6.45 | 6.18 | 11.06 | 7.36 | 1.099114589 | 6.85E-09 | 1.93E-07 | Up in group#2 |
| Fgf14 | 0.29 | 0.21 | 0.76 | 0.23 | 1.23 | 0.81 | 0.46 | 0.61 | 1.098690739 | 0.005120413 | 0.022350274 | Up in group#2 |
| Alox5ap | 6.78 | 5.38 | 8.95 | 7.62 | 12.28 | 11.05 | 20.52 | 15.55 | 1.098660712 | 1.32E-08 | 3.43E-07 | Up in group#2 |
| Bst1 | 0.47 | 0.45 | 0.94 | 0.6 | 1 | 0.95 | 1.8 | 1.35 | 1.09865452 | 9.32E-05 | 0.000814845 | Up in group#2 |
| Iqgap2 | 1.07 | 0.91 | 1.6 | 1.19 | 2.12 | 1.65 | 3.3 | 2.79 | 1.097738754 | 5.27E-08 | 1.17E-06 | Up in group#2 |
| Lilrb4a | 5.59 | 4.31 | 6.38 | 5.46 | 8.67 | 8.39 | 15.74 | 12.08 | 1.097027923 | 6.02E-09 | 1.71E-07 | Up in group#2 |
| Tlr8 | 0.56 | 0.39 | 0.86 | 0.66 | 0.84 | 0.85 | 1.76 | 1.49 | 1.092689074 | 2.57E-05 | 0.000272752 | Up in group#2 |
| Pcdh9 | 0.37 | 0.3 | 0.25 | 0.29 | 0.59 | 0.5 | 0.66 | 0.7 | 1.09237197 | 1.01E-07 | 2.13E-06 | Up in group#2 |
| Col5a3 | 5.59 | 4.82 | 4.87 | 4.43 | 8.04 | 8.5 | 12.08 | 12.05 | 1.091351357 | 9.23E-17 | 1.23E-14 | Up in group#2 |
| AV026068 | 1.73 | 1.45 | 2.58 | 1.91 | 3.69 | 3.66 | 4.72 | 3.65 | 1.089255331 | 4.98E-06 | 6.53E-05 | Up in group#2 |
| Gfra1 | 1.77 | 1.78 | 1.45 | 1.6 | 2.58 | 2.74 | 4.06 | 4.18 | 1.086440037 | 7.41E-11 | 3.39E-09 | Up in group#2 |
| Pknox2 | 1.16 | 0.98 | 1.27 | 1.01 | 1.46 | 1.83 | 2.15 | 3.61 | 1.082904991 | 5.40E-06 | 6.98E-05 | Up in group#2 |
| Lin7a | 0.91 | 0.9 | 0.9 | 0.86 | 1.37 | 1.34 | 2.29 | 2.31 | 1.081900026 | 1.12E-08 | 2.99E-07 | Up in group#2 |
| Ccl11 | 1.2 | 1.2 | 1.09 | 1.18 | 2.55 | 1.94 | 2.39 | 2.7 | 1.078894041 | 4.59E-06 | 6.08E-05 | Up in group#2 |
| Mdga2 | 0.06 | 0.04 | 0.09 | 0.08 | 0.1 | 0.13 | 0.19 | 0.14 | 1.078881491 | 0.000632979 | 0.004095911 | Up in group#2 |
| Hk3 | 0.39 | 0.35 | 0.92 | 0.37 | 0.76 | 0.48 | 1.65 | 1.25 | 1.076220652 | 0.003363644 | 0.016003449 | Up in group#2 |
| Baiap3 | 0.11 | 0.09 | 0.09 | 0.1 | 0.19 | 0.13 | 0.31 | 0.17 | 1.073088803 | 0.004068233 | 0.018706875 | Up in group#2 |
| Wnt9b | 0.88 | 0.75 | 1.28 | 0.86 | 2.06 | 1.57 | 1.91 | 2.14 | 1.072834156 | 4.90E-10 | 1.84E-08 | Up in group#2 |
| Emilin2 | 1.73 | 1.72 | 2.7 | 2.24 | 3.75 | 2.69 | 5.53 | 5.06 | 1.071553762 | 6.92E-08 | 1.50E-06 | Up in group#2 |
| Tlr7 | 1.19 | 1.04 | 1.67 | 1.24 | 1.97 | 1.83 | 3.62 | 3.02 | 1.070193526 | 6.56E-07 | 1.09E-05 | Up in group#2 |
| Col23a1 | 0.45 | 0.61 | 1.48 | 0.5 | 1.8 | 1.6 | 1.52 | 1.25 | 1.070086386 | 0.000162437 | 0.001315 | Up in group#2 |
| Eps8 | 6.56 | 5.91 | 7.03 | 7.08 | 10.22 | 14.64 | 14.41 | 15.31 | 1.067466249 | 2.79E-19 | 6.88E-17 | Up in group#2 |
| Svopl | 0.37 | 0.23 | 0.34 | 0.45 | 0.7 | 0.52 | 0.89 | 0.72 | 1.062121695 | 0.000254509 | 0.001914854 | Up in group#2 |
| 2810468N07I | 0.54 | 0.4 | 0.32 | 0.34 | 0.74 | 0.83 | 0.89 | 0.77 | 1.061340854 | 0.001343903 | 0.007595965 | Up in group#2 |

| | | | | | | | | | | | |
|---|---|---|---|---|---|---|---|---|---|---|---|
| Klf5 | 1 | 1.2 | 0.92 | 0.87 | 1.34 | 1.98 | 2.14 | 2.58 | 1.056278103 | 1.13E-05 | 0.000132274 | Up in group#2 |
| Vtn | 32.14 | 28.81 | 27.06 | 28.46 | 46.44 | 53.06 | 65.7 | 68.41 | 1.054159966 | 1.64E-20 | 4.85E-18 | Up in group#2 |
| Plp1 | 1.81 | 2.13 | 2.11 | 2.45 | 3.25 | 3.32 | 5.03 | 5.41 | 1.052728547 | 1.19E-09 | 4.11E-08 | Up in group#2 |
| Erbb3 | 0.44 | 0.45 | 0.41 | 0.4 | 0.56 | 0.54 | 1.04 | 1.24 | 1.051633488 | 0.000139191 | 0.001149165 | Up in group#2 |
| Greb1l | 0.21 | 0.24 | 0.53 | 0.26 | 0.71 | 0.52 | 0.59 | 0.68 | 1.049392307 | 1.47E-05 | 0.000168959 | Up in group#2 |
| Cebpd | 39.67 | 36.29 | 37.15 | 35.94 | 60.45 | 71.3 | 81.1 | 84.96 | 1.048558798 | 2.44E-26 | 1.48E-23 | Up in group#2 |
| Rgs18 | 0.65 | 0.35 | 0.75 | 0.73 | 1.41 | 0.73 | 1.57 | 1.28 | 1.045386438 | 0.000261752 | 0.001964913 | Up in group#2 |
| Aldh1a1 | 4.32 | 3.99 | 5.31 | 4.54 | 7.98 | 6.35 | 12.93 | 8.91 | 1.042085635 | 5.57E-09 | 1.61E-07 | Up in group#2 |
| Csdc2 | 1.04 | 1.45 | 1.79 | 1.18 | 2.22 | 2.68 | 3.13 | 2.85 | 1.041387212 | 2.96E-08 | 7.02E-07 | Up in group#2 |
| C5ar2 | 0.46 | 0.43 | 0.71 | 0.2 | 0.88 | 0.51 | 1.28 | 0.91 | 1.041350354 | 0.001221529 | 0.007035946 | Up in group#2 |
| Tbxas1 | 1.23 | 0.75 | 1.46 | 1.11 | 1.82 | 1.61 | 3.47 | 2.16 | 1.041175229 | 6.68E-05 | 0.000610917 | Up in group#2 |
| Cd300lb | 0.35 | 0.24 | 0.56 | 0.37 | 0.66 | 0.39 | 0.94 | 1.06 | 1.040536215 | 0.001806613 | 0.009650489 | Up in group#2 |
| Pcolce2 | 2.95 | 2.76 | 3.11 | 2.51 | 4.19 | 4.57 | 8.28 | 5.48 | 1.040420753 | 1.82E-07 | 3.56E-06 | Up in group#2 |
| Syt17 | 0.3 | 0.49 | 0.32 | 0.28 | 0.44 | 0.44 | 1.1 | 0.8 | 1.040263267 | 0.007975066 | 0.031807116 | Up in group#2 |
| Pla1a | 4.33 | 4.61 | 4.38 | 3.8 | 6.23 | 7.05 | 11.31 | 9.42 | 1.038810754 | 1.37E-09 | 4.57E-08 | Up in group#2 |
| Vcan | 1.82 | 1.44 | 1.8 | 1.31 | 2.57 | 2.11 | 4.33 | 3.6 | 1.037421232 | 3.64E-09 | 1.09E-07 | Up in group#2 |
| Dcn | 146.84 | 166.03 | 192.19 | 133.19 | 240.15 | 243.69 | 414.49 | 372.06 | 1.03679252 | 1.26E-10 | 5.39E-09 | Up in group#2 |
| Irs2 | 6.68 | 5.59 | 5.76 | 4.95 | 8.42 | 9.36 | 14.06 | 13.78 | 1.036475812 | 1.27E-11 | 6.89E-10 | Up in group#2 |
| Bmp2 | 2.04 | 1.64 | 1.6 | 1.42 | 2.65 | 2.63 | 4.04 | 3.99 | 1.036338977 | 5.81E-09 | 1.66E-07 | Up in group#2 |
| Amy1 | 0.68 | 0.63 | 1.12 | 0.71 | 1.52 | 1.47 | 1.46 | 1.68 | 1.032095267 | 1.05E-05 | 0.000124547 | Up in group#2 |
| Pkp4 | 116.53 | 108.85 | 121.98 | 140.44 | 205.2 | 269.53 | 257.86 | 230.78 | 1.03136159 | 2.36E-22 | 9.51E-20 | Up in group#2 |
| Ang | 2.38 | 3.18 | 3.13 | 3.21 | 4.51 | 4.49 | 6.33 | 8.11 | 1.03093531 | 6.07E-06 | 7.75E-05 | Up in group#2 |
| Ltc4s | 3.61 | 1.77 | 5.17 | 2.13 | 5.56 | 5.01 | 7.72 | 5.98 | 1.03041831 | 0.000301685 | 0.002215961 | Up in group#2 |
| Ssc5d | 1.07 | 0.93 | 0.73 | 0.52 | 1.05 | 1.35 | 2.14 | 1.89 | 1.029823268 | 2.55E-05 | 0.000270849 | Up in group#2 |
| Efemp1 | 1.22 | 1.45 | 4.33 | 1.72 | 3.57 | 3.16 | 6.11 | 4.33 | 1.028042541 | 0.001152118 | 0.006735188 | Up in group#2 |
| Ptgs2 | 8.79 | 8.01 | 17.48 | 5.99 | 17.09 | 18.32 | 24.94 | 18.94 | 1.027676253 | 7.14E-06 | 8.88E-05 | Up in group#2 |
| Sfmbt2 | 0.13 | 0.07 | 0.07 | 0.09 | 0.16 | 0.16 | 0.23 | 0.14 | 1.02731461 | 0.001132881 | 0.006654864 | Up in group#2 |
| Ggt5 | 2.08 | 1.85 | 1.78 | 1.9 | 2.91 | 3.62 | 4.34 | 4.08 | 1.027276081 | 2.39E-13 | 1.78E-11 | Up in group#2 |
| Abcc9 | 31.24 | 29.52 | 25.49 | 27.59 | 42.4 | 45.14 | 69.06 | 67.86 | 1.025731113 | 5.13E-13 | 3.50E-11 | Up in group#2 |
| Fgl2 | 10.17 | 9.04 | 8.97 | 8.09 | 13.64 | 14.21 | 23.09 | 20.46 | 1.024122082 | 2.00E-12 | 1.26E-10 | Up in group#2 |
| Scara5 | 2.06 | 2.07 | 1.91 | 1.78 | 2.23 | 2.74 | 5.67 | 4.77 | 1.023671455 | 8.97E-06 | 0.000108574 | Up in group#2 |
| Nova1 | 0.22 | 0.25 | 0.43 | 0.3 | 0.39 | 0.51 | 0.86 | 0.61 | 1.02268607 | 7.74E-05 | 0.000692515 | Up in group#2 |
| Sorcs1 | 0.17 | 0.2 | 0.18 | 0.2 | 0.32 | 0.41 | 0.43 | 0.32 | 1.022646615 | 7.08E-06 | 8.83E-05 | Up in group#2 |
| Shc2 | 0.75 | 0.83 | 0.78 | 0.54 | 1.39 | 1.25 | 1.77 | 1.31 | 1.021202757 | 4.13E-08 | 9.43E-07 | Up in group#2 |
| Tlr13 | 0.78 | 0.75 | 0.79 | 0.71 | 1.23 | 0.84 | 2.32 | 1.57 | 1.021048827 | 4.07E-05 | 0.000401358 | Up in group#2 |
| Slc41a2 | 0.83 | 0.65 | 0.94 | 0.56 | 1.33 | 1.28 | 1.63 | 1.63 | 1.02063762 | 2.16E-08 | 5.31E-07 | Up in group#2 |
| Ptprd | 0.2 | 0.31 | 0.27 | 0.25 | 0.46 | 0.41 | 0.59 | 0.55 | 1.019475267 | 4.34E-07 | 7.64E-06 | Up in group#2 |
| Rgs5 | 667.8 | 612.71 | 581.2 | 642.01 | 1082.46 | 1073.99 | 1367.85 | 1384.9 | 1.019000923 | 3.92E-20 | 1.06E-17 | Up in group#2 |
| Carmil1 | 0.87 | 0.74 | 1.17 | 1 | 1.6 | 1.48 | 2.1 | 2.24 | 1.018438738 | 5.63E-09 | 1.62E-07 | Up in group#2 |
| Pamr1 | 0.45 | 0.54 | 0.96 | 0.61 | 1.02 | 1.03 | 1.48 | 1.48 | 1.018163125 | 3.22E-05 | 0.000328769 | Up in group#2 |
| Bcl2l1 | 37.41 | 28.84 | 35.81 | 33.74 | 46.47 | 79.33 | 73.11 | 66.77 | 1.016099449 | 8.32E-14 | 6.64E-12 | Up in group#2 |
| Cacnb4 | 0.36 | 0.39 | 0.33 | 0.24 | 0.37 | 0.52 | 0.84 | 0.79 | 1.01569841 | 6.90E-05 | 0.000628967 | Up in group#2 |

| | | | | | | | | | | | | |
|---|---|---|---|---|---|---|---|---|---|---|---|---|
| Serpinb8 | 0.91 | 1.02 | 1.27 | 0.84 | 1.83 | 1.67 | 2.7 | 2.69 | 1.013063049 | 1.75E-06 | 2.62E-05 | Up in group#2 |
| Tmem37 | 0.87 | 0.78 | 1.33 | 0.84 | 1.28 | 1.57 | 2.77 | 1.85 | 1.009719461 | 0.001431751 | 0.008007826 | Up in group#2 |
| Vasn | 5.66 | 5.13 | 4.59 | 4.05 | 7.66 | 8.05 | 11.53 | 10.64 | 1.009558439 | 2.26E-12 | 1.41E-10 | Up in group#2 |
| Rtn4rl2 | 2.42 | 1.98 | 2.26 | 1.94 | 2.77 | 4.47 | 5.27 | 4.17 | 1.007856318 | 2.63E-06 | 3.75E-05 | Up in group#2 |
| Abca9 | 3.8 | 2.94 | 3.6 | 3.23 | 4.82 | 4.9 | 9.43 | 7.23 | 1.006459112 | 7.98E-09 | 2.22E-07 | Up in group#2 |
| Qpct | 1.49 | 1.85 | 1.93 | 1.83 | 3 | 2.97 | 4.21 | 3.71 | 1.006085478 | 7.31E-09 | 2.04E-07 | Up in group#2 |
| Jdp2 | 29.5 | 21.73 | 22.81 | 24.13 | 37.82 | 51.88 | 52.21 | 48.11 | 1.004841748 | 2.24E-17 | 3.55E-15 | Up in group#2 |
| Adamts8 | 0.52 | 0.4 | 0.89 | 0.51 | 1.56 | 0.96 | 1.02 | 1 | 1.003401961 | 1.74E-05 | 0.000196308 | Up in group#2 |
| Lrrc23 | 0.5 | 0.33 | 0.47 | 0.36 | 0.53 | 0.63 | 0.97 | 1.09 | 1.001236688 | 0.004452549 | 0.020058447 | Up in group#2 |
| Pou5f2 | 0.48 | 0.54 | 0.53 | 0.99 | 0.32 | 0.35 | 0.15 | 0.39 | -1.002909956 | 0.013040522 | 0.046677674 | Up in group#1 |
| Gm4841 | 1.23 | 0.85 | 2.52 | 0.97 | 0.7 | 0.83 | 0.49 | 0.64 | -1.006641858 | 0.001081453 | 0.006437174 | Up in group#1 |
| Ccnb1 | 0.88 | 0.33 | 0.69 | 0.54 | 0.29 | 0.35 | 0.25 | 0.26 | -1.007928004 | 0.003231013 | 0.015516552 | Up in group#1 |
| Kif22 | 2.4 | 0.99 | 1.44 | 1.06 | 0.66 | 0.91 | 0.65 | 0.57 | -1.02278721 | 0.000630651 | 0.004084816 | Up in group#1 |
| Ifit3b | 17.06 | 18.9 | 21.87 | 12.25 | 7.41 | 9.48 | 5.42 | 10.89 | -1.026714197 | 8.87E-08 | 1.88E-06 | Up in group#1 |
| Lrp11 | 1.25 | 1.01 | 1.32 | 1.14 | 0.52 | 0.56 | 0.58 | 0.62 | -1.031695985 | 1.51E-07 | 3.02E-06 | Up in group#1 |
| Aldh1l1 | 9.08 | 8.25 | 10.47 | 4.46 | 4.86 | 3.46 | 3.74 | 3.07 | -1.043358916 | 2.53E-07 | 4.77E-06 | Up in group#1 |
| Ccr3 | 0.31 | 0.3 | 0.22 | 0.5 | 0.22 | 0.08 | 0.15 | 0.18 | -1.044650372 | 0.004891614 | 0.021562935 | Up in group#1 |
| Dtl | 0.38 | 0.27 | 0.46 | 0.33 | 0.11 | 0.22 | 0.21 | 0.14 | -1.045359143 | 0.000690963 | 0.004396347 | Up in group#1 |
| Orc1 | 0.47 | 0.43 | 0.44 | 0.23 | 0.12 | 0.25 | 0.16 | 0.2 | -1.046235589 | 0.002885906 | 0.014160507 | Up in group#1 |
| Sult5a1 | 2.48 | 1.67 | 2.97 | 1.69 | 1.14 | 0.52 | 1.3 | 1.19 | -1.046656117 | 0.000445881 | 0.003070215 | Up in group#1 |
| Ifit2 | 59.44 | 61.88 | 68.22 | 37.29 | 21.93 | 30.77 | 15.83 | 37.12 | -1.048097418 | 7.75E-07 | 1.27E-05 | Up in group#1 |
| Slco4a1 | 0.43 | 0.17 | 0.2 | 0.25 | 0.19 | 0.1 | 0.12 | 0.08 | -1.062139295 | 0.014104707 | 0.049481662 | Up in group#1 |
| Slc16a4 | 2.62 | 1.92 | 2.58 | 2.15 | 1.14 | 1.19 | 1.14 | 0.77 | -1.069643192 | 5.65E-08 | 1.25E-06 | Up in group#1 |
| Dsc2 | 0.57 | 0.63 | 1.3 | 0.52 | 0.37 | 0.36 | 0.28 | 0.38 | -1.072916668 | 0.000158823 | 0.001288878 | Up in group#1 |
| Cdkn1c | 139.89 | 139.16 | 157.65 | 147.91 | 78.79 | 53.74 | 68.6 | 67.54 | -1.074244195 | 6.07E-32 | 6.74E-29 | Up in group#1 |
| Apln | 38.12 | 34.88 | 28.25 | 29.99 | 17.93 | 14.35 | 14.01 | 14.09 | -1.076238019 | 1.92E-24 | 9.14E-22 | Up in group#1 |
| Gm7609 | 13.68 | 12.29 | 20.57 | 11.13 | 6.52 | 6.78 | 5.09 | 7.93 | -1.07974989 | 1.21E-09 | 4.16E-08 | Up in group#1 |
| Gm4951 | 5.96 | 4.26 | 9.58 | 3.94 | 3.18 | 3.81 | 1.5 | 2.28 | -1.086798227 | 8.06E-05 | 0.000716218 | Up in group#1 |
| Gm10432 | 0.2 | 0.24 | 0.25 | 0.18 | 0.12 | 0.08 | 0.12 | 0.07 | -1.088459048 | 0.00104376 | 0.006258331 | Up in group#1 |
| Melk | 0.63 | 0.37 | 0.52 | 0.4 | 0.37 | 0.29 | 0.13 | 0.08 | -1.103476158 | 0.00397828 | 0.018344445 | Up in group#1 |
| Sgpp2 | 0.5 | 0.42 | 0.42 | 0.49 | 0.24 | 0.12 | 0.26 | 0.21 | -1.10422058 | 4.40E-05 | 0.000428177 | Up in group#1 |
| Oas1a | 17.39 | 14.79 | 17.96 | 13.16 | 7.11 | 7.56 | 5.21 | 8.52 | -1.106247139 | 7.21E-15 | 7.27E-13 | Up in group#1 |
| Halr1 | 7.32 | 6.33 | 12.32 | 9.24 | 5.96 | 2.64 | 4.52 | 2.69 | -1.10842647 | 7.09E-05 | 0.000643043 | Up in group#1 |
| Gbp11 | 0.49 | 0.37 | 0.58 | 0.24 | 0.15 | 0.26 | 0.11 | 0.22 | -1.108737318 | 0.003573166 | 0.016879692 | Up in group#1 |
| Oasl2 | 36.67 | 35.09 | 45.9 | 27.33 | 17.46 | 15.95 | 13.14 | 18.36 | -1.111476003 | 2.64E-17 | 4.09E-15 | Up in group#1 |
| Ifi44 | 28.94 | 24.3 | 31.44 | 21.54 | 13.4 | 11.7 | 8.02 | 13.48 | -1.141516206 | 3.24E-15 | 3.40E-13 | Up in group#1 |
| Smco3 | 0.44 | 0.6 | 0.49 | 0.51 | 0.18 | 0.38 | 0.17 | 0.14 | -1.155903509 | 0.001839714 | 0.009811564 | Up in group#1 |
| Hcn2 | 0.7 | 0.98 | 1.03 | 0.49 | 0.36 | 0.37 | 0.36 | 0.3 | -1.156091369 | 2.47E-05 | 0.000264504 | Up in group#1 |
| Scn3b | 2.18 | 1.82 | 2.33 | 1.81 | 0.9 | 0.67 | 0.97 | 0.97 | -1.156819033 | 1.47E-12 | 9.42E-11 | Up in group#1 |
| Gkn3 | 1.44 | 1.69 | 2.74 | 1 | 0.96 | 0.62 | 0.97 | 0.42 | -1.162426496 | 0.003146053 | 0.01516885 | Up in group#1 |
| Spin2c | 1.72 | 1.57 | 1.6 | 1.11 | 1.11 | 0.56 | 0.4 | 0.53 | -1.164744938 | 0.000190159 | 0.001495773 | Up in group#1 |
| Acta1 | 18.28 | 15.42 | 19.4 | 14.34 | 9.37 | 7.33 | 7.13 | 5.31 | -1.166135691 | 3.86E-15 | 4.02E-13 | Up in group#1 |

| | | | | | | | | | | | | |
|---|---|---|---|---|---|---|---|---|---|---|---|---|
| Lrmda | 1.41 | 2.05 | 1.84 | 1.73 | 0.87 | 0.75 | 0.87 | 0.53 | -1.166334379 | 0.000132531 | 0.001097587 | Up in group#1 |
| Ifit3 | 58.27 | 63.24 | 74.88 | 38.89 | 24.55 | 27.37 | 16.88 | 32.39 | -1.168647996 | 2.27E-10 | 9.11E-09 | Up in group#1 |
| Gm12216 | 3.4 | 4.61 | 4.66 | 4.03 | 2.34 | 1.74 | 1.21 | 1.9 | -1.169030689 | 3.20E-08 | 7.55E-07 | Up in group#1 |
| Cxcl9 | 70.61 | 65.19 | 60.66 | 44.78 | 22.49 | 36.32 | 14.65 | 29.69 | -1.175031488 | 1.44E-08 | 3.72E-07 | Up in group#1 |
| Prtg | 0.12 | 0.13 | 0.24 | 0.12 | 0.07 | 0.03 | 0.08 | 0.07 | -1.195120136 | 0.000678035 | 0.004330033 | Up in group#1 |
| Camk2a | 1.61 | 1.94 | 1.66 | 1.52 | 0.8 | 0.62 | 0.75 | 0.65 | -1.209511205 | 1.44E-13 | 1.10E-11 | Up in group#1 |
| Map6 | 11.23 | 14.48 | 13.85 | 14.81 | 5.81 | 6.88 | 4.77 | 4.76 | -1.225444496 | 8.33E-21 | 2.77E-18 | Up in group#1 |
| Dlk2 | 1.06 | 0.75 | 1.08 | 0.69 | 0.39 | 0.58 | 0.33 | 0.17 | -1.231633099 | 0.000676275 | 0.004325626 | Up in group#1 |
| Nppb | 5.89 | 6.1 | 4.97 | 5.84 | 2.2 | 1.49 | 2.88 | 2.67 | -1.250541902 | 1.64E-08 | 4.17E-07 | Up in group#1 |
| Gm5431 | 4.66 | 2.89 | 2.76 | 1.64 | 1.05 | 1.69 | 0.84 | 1.25 | -1.254978052 | 5.07E-06 | 6.62E-05 | Up in group#1 |
| Dhx58 | 7.44 | 5.78 | 7.63 | 5.17 | 2.98 | 2.47 | 2.12 | 2.89 | -1.265956085 | 5.94E-16 | 6.93E-14 | Up in group#1 |
| F2rl3 | 2.92 | 2.2 | 2.68 | 2.22 | 1.06 | 1.06 | 1.09 | 0.8 | -1.270772902 | 5.84E-11 | 2.69E-09 | Up in group#1 |
| BC051537 | 0.58 | 0.44 | 0.37 | 0.74 | 0.32 | 0.25 | 0.21 | 0.06 | -1.273252471 | 0.002817394 | 0.013890866 | Up in group#1 |
| Exo1 | 0.24 | 0.17 | 0.19 | 0.2 | 0.12 | 0.07 | 0.08 | 0.05 | -1.281084191 | 0.000171934 | 0.001378475 | Up in group#1 |
| Parpbp | 0.24 | 0.26 | 0.19 | 0.37 | 0.15 | 0.08 | 0.08 | 0.1 | -1.287520058 | 0.000702737 | 0.004463545 | Up in group#1 |
| A930017M01 | 0.35 | 0.23 | 0.29 | 0.42 | 0.15 | 0.12 | 0.13 | 0.11 | -1.297181487 | 0.000185841 | 0.001469626 | Up in group#1 |
| Rarres1 | 5.86 | 6.13 | 5.96 | 5.85 | 2.62 | 1.91 | 2.13 | 2.47 | -1.333197022 | 5.38E-18 | 9.95E-16 | Up in group#1 |
| Ifi213 | 2.7 | 2.37 | 2.96 | 2.12 | 1.19 | 1.26 | 0.49 | 0.91 | -1.348511487 | 2.14E-09 | 6.72E-08 | Up in group#1 |
| Slitrk2 | 0.18 | 0.26 | 0.33 | 0.27 | 0.15 | 0.04 | 0.11 | 0.1 | -1.349517072 | 2.52E-05 | 0.000268673 | Up in group#1 |
| Fut4 | 1.28 | 0.9 | 0.95 | 1.18 | 0.58 | 0.3 | 0.49 | 0.26 | -1.369265215 | 1.82E-08 | 4.57E-07 | Up in group#1 |
| Epb41l4a | 9.45 | 9.64 | 10.7 | 9.06 | 4.82 | 3.2 | 2.98 | 3.59 | -1.369488827 | 2.08E-29 | 2.13E-26 | Up in group#1 |
| Ticrr | 0.14 | 0.11 | 0.07 | 0.04 | 0.03 | 0.02 | 0.05 | 0.02 | -1.390233632 | 0.003719503 | 0.017431754 | Up in group#1 |
| Rtkn2 | 0.44 | 0.43 | 0.42 | 0.6 | 0.19 | 0.15 | 0.19 | 0.16 | -1.400435048 | 2.61E-07 | 4.91E-06 | Up in group#1 |
| Zfp941 | 0.42 | 0.61 | 0.27 | 0.24 | 0.19 | 0.09 | 0.1 | 0.18 | -1.416164115 | 0.000209168 | 0.001627037 | Up in group#1 |
| 5830444B04F | 3.16 | 3.17 | 4.36 | 3.27 | 1.03 | 1.85 | 1.15 | 1.58 | -1.42636331 | 9.23E-13 | 6.00E-11 | Up in group#1 |
| Rec8 | 0.95 | 0.54 | 0.33 | 0.32 | 0.14 | 0.24 | 0.18 | 0.24 | -1.441190496 | 0.000961408 | 0.005840816 | Up in group#1 |
| Adamts17 | 0.5 | 0.49 | 0.4 | 0.27 | 0.15 | 0.11 | 0.16 | 0.17 | -1.445122297 | 1.83E-07 | 3.57E-06 | Up in group#1 |
| Cfap74 | 1.62 | 1.15 | 1.76 | 1.05 | 0.42 | 0.47 | 0.34 | 0.74 | -1.445435401 | 1.84E-08 | 4.60E-07 | Up in group#1 |
| Il4i1 | 2.25 | 1.52 | 1.55 | 1.36 | 0.9 | 0.58 | 0.46 | 0.44 | -1.447244367 | 3.64E-08 | 8.45E-07 | Up in group#1 |
| Apol10b | 72.25 | 102.94 | 88.52 | 73.93 | 34.34 | 35.16 | 19.87 | 29.81 | -1.454662402 | 1.64E-21 | 5.90E-19 | Up in group#1 |
| Muc13 | 1.52 | 0.96 | 0.84 | 0.57 | 0.33 | 0.28 | 0.4 | 0.35 | -1.460302406 | 1.50E-06 | 2.27E-05 | Up in group#1 |
| Nrep | 73.24 | 83.41 | 98.75 | 81.03 | 36.88 | 30.73 | 26.98 | 22.5 | -1.470421372 | 2.60E-33 | 3.15E-30 | Up in group#1 |
| Oas2 | 41.56 | 31.23 | 42.38 | 33.14 | 15.08 | 11.63 | 10.17 | 14.59 | -1.481994389 | 1.27E-33 | 1.87E-30 | Up in group#1 |
| Cd247 | 6.93 | 9.78 | 9.31 | 7.29 | 3.6 | 2.91 | 2.77 | 2.26 | -1.499569647 | 2.73E-20 | 7.58E-18 | Up in group#1 |
| Apol9b | 1.18 | 0.42 | 1.4 | 0.3 | 0.35 | 0.26 | 0.17 | 0.33 | -1.505615339 | 0.001716578 | 0.009285 | Up in group#1 |
| Scgb3a1 | 8.47 | 10.49 | 14.38 | 7.47 | 5.48 | 3.09 | 2.27 | 2.61 | -1.526898504 | 8.11E-08 | 1.73E-06 | Up in group#1 |
| Akap5 | 13.38 | 13.67 | 10.68 | 9.73 | 5.97 | 3.92 | 3.32 | 2.69 | -1.537107419 | 3.86E-17 | 5.71E-15 | Up in group#1 |
| Rnaset2b | 4.6 | 5.19 | 2.6 | 5.59 | 0.89 | 0 | 2.55 | 2.51 | -1.543302113 | 0.00628175 | 0.026306308 | Up in group#1 |
| Ido1 | 3.95 | 4.58 | 5 | 3.57 | 1.38 | 1.34 | 1.43 | 1.33 | -1.5670905 | 2.78E-18 | 5.37E-16 | Up in group#1 |
| B230344G16I | 0.79 | 0.44 | 0.36 | 0.1 | 0.2 | 0.13 | 0.07 | 0.14 | -1.593082104 | 0.002358611 | 0.012014716 | Up in group#1 |
| Zbp1 | 12.01 | 8.8 | 13.12 | 5.88 | 2.89 | 3.83 | 1.47 | 2.6 | -1.626054839 | 1.80E-08 | 4.55E-07 | Up in group#1 |
| Clca2 | 0.79 | 0.49 | 0.57 | 0.48 | 0.21 | 0.2 | 0.12 | 0.19 | -1.638266718 | 1.24E-08 | 3.27E-07 | Up in group#1 |

| | | | | | | | | | | | |
|---|---|---|---|---|---|---|---|---|---|---|---|
| Prnd | 33.14 | 25.92 | 24.32 | 21.57 | 11.13 | 7.75 | 8.05 | 5.66 | -1.675956651 | 3.30E-25 | 1.69E-22 | Up in group#1 |
| Pimreg | 2.41 | 1.19 | 1.25 | 1.13 | 0.45 | 0.56 | 0.35 | 0.43 | -1.678449311 | 5.27E-07 | 9.02E-06 | Up in group#1 |
| Lrrc55 | 0.74 | 0.82 | 0.66 | 0.55 | 0.33 | 0.09 | 0.25 | 0.16 | -1.700904117 | 1.15E-07 | 2.38E-06 | Up in group#1 |
| Has3 | 3.95 | 3.33 | 4.1 | 2.76 | 1.39 | 1.08 | 0.8 | 0.8 | -1.751774791 | 5.58E-24 | 2.56E-21 | Up in group#1 |
| Oas1g | 3.05 | 2.05 | 3.1 | 1.79 | 0.9 | 0.52 | 0.58 | 0.77 | -1.802252045 | 2.27E-13 | 1.71E-11 | Up in group#1 |
| Col17a1 | 0.21 | 0.12 | 0.14 | 0.15 | 0.05 | 0.04 | 0.03 | 0.04 | -1.834285022 | 8.99E-06 | 0.000108751 | Up in group#1 |
| Lrrn1 | 5.43 | 4.23 | 5.34 | 3.84 | 1.48 | 1.04 | 0.83 | 1.39 | -1.943751386 | 3.04E-29 | 2.89E-26 | Up in group#1 |
| Cacna1b | 0.32 | 0.3 | 0.21 | 0.25 | 0.09 | 0.06 | 0.07 | 0.05 | -1.995883226 | 2.68E-13 | 1.95E-11 | Up in group#1 |
| Aplnr | 83.67 | 88.1 | 92.3 | 80.84 | 26.84 | 17.54 | 16.8 | 13.11 | -2.17157951 | 1.28E-47 | 3.42E-44 | Up in group#1 |
| Apol9a | 0.68 | 0.4 | 1.1 | 0.42 | 0.19 | 0.16 | 0.03 | 0.1 | -2.363042456 | 3.03E-06 | 4.23E-05 | Up in group#1 |
| Arsi | 4.73 | 4.67 | 6.51 | 4.56 | 1.28 | 0.38 | 0.76 | 0.56 | -2.748431484 | 7.09E-29 | 5.90E-26 | Up in group#1 |
| Bmp10 | 0 | 0 | 1.41 | 0.31 | 0.08 | 0 | 0 | 0.11 | -2.957126457 | 0.010157401 | 0.038552121 | Up in group#1 |
| 4933417O13I | 0.57 | 0.76 | 0.68 | 0.56 | 0.04 | 0.06 | 0.1 | 0.06 | -3.160451523 | 5.47E-11 | 2.55E-09 | Up in group#1 |
| Hmga1b | 0.79 | 0.05 | 0 | 0.05 | 0 | 0 | 0 | 0 | -5.770384151 | 5.65E-05 | 0.000527923 | Up in group#1 |
