## Supplementary data for "Mast cells participate in the development of diastolic dysfunction in diabetic obese mice"

**Supplemental table**

| | | |
|---|---|---|
| 18S | F | 5′-CGCGGTTCTATTTTGTTGGT-3′ |
| | R | 5′-AGTCGGCATCGTTTATGGTC-3′ |
| Myh7 | F | 5′-GGATGACGTCACCTCCAACA-3′ |
| | R | 5′-AGATCAGAGCCTCCTTCTCGT-3′ |
| Vcam1 | F | 5′-CGTACACCATCCGCCAGGCA-3′ |
| | R | 5′-TAGAGTGCAAGGAGTTCGGGCG-3′ |
| Icam1 | F | 5′-TGGCCTGGGGGATGCACACT-3′ |
| | R | 5′-CCACCGGGCTGTAGGTGGGT-3′ |
| Sele | F | 5′-ACGTCCCGGGAAAGATGAAC-3′ |
| | R | 5′-GTCAGGAGTGAGGTTCCTGC-3′ |
| Ccl7 | F | 5′-AAGTGGGTCGAGGAGGCTAT-3′ |
| | R | 5′-AGCTCCTATCCCTTAGGACCG-3′ |
| Il6 | F | 5′-CACTTCACAAGTCGGAGGCT-3′ |
| | R | 5′-CTGCAAGTGCATCATCGTTGT-3′ |
| Cpa3 | F | 5′-GCCCTTGTTTTGAAACGTGCT-3′ |
| | R | 5′-TTAAAGTGGGGCTGTTGGGAG-3′ |
| Fcer1a | F | 5′-TTCTCCACTGTCAAAGGCCA-3′ |
| | R | 5′-GGCAGTGTTTATTGAGTATTTGCTA-3′ |
| Tpsab1 | F | 5′-CTTGGACTGGATCCACCACT-3′ |
| | R | 5′-TTGAGGCATAGCAGAGAGCG-3′ |
| Cma1 | F | 5′-CAGCCTGTGAGGAAATCTGGAA-3′ |
| | R | 5′-GCAGTTGACAATCTGGGTCTTTA-3′ |
| Alox5 | F | 5′-CATACCACATGCTGAGGTCCA-3′ |
| | R | 5′-CTACAAAAGCAGAAAGGGCCAC-3′ |
| ANP | F | 5′-CGTCTTGGCCTTTTGGCTTC-3′ |
| | R | 5′-GGTGGTCTAGCAGGTTCTTGAAA-3′ |
| BNP | F | 5′-AAGCTGCTGGAGCTGATAAGA-3′ |
| | R | 5′-GTTACAGCCCAAACGACTGAC-3′ |
| Col1a1 | F | 5′-CAACCTCAAGAAGGCCCTGC-3′ |
| | R | 5′-TGTCCAAGGGAGCCACATCG-3′ |
| Col3a1 | F | 5′-AGCACGAGGTCTTGCTGGAC-3′ |
| | R | 5′-ACCAGCTGTACCAGGCTGAC-3′ |
| Lox | F | 5′-CGCTAGGCACTCTTTGTAACAG-3′ |
| | R | 5′-AGTAGTTCATGACTAAGGGCTCA-3′ |
| Titin isoform N2-A | F | 5′-GAGACATTGCTCCGCTTTTC-3′ |
| | R | 5′-GATCTCCAAAGAGGCTGTC-3′ |
| Titin isoform N2-B | F | 5′- ACAGTGGGAAAGCAAAGACATC-3′ |
| | R | 5′- AGGTGGCCCAGAGCTACTTC-3′ |
| MEF2C | F | 5′- TAAGCAGGCAAGGGTCACTG-3′ |
| | R | 5′- AAAGTCCAGCTTATGCCGCT-3′ |
| Cx43 | F | 5′- ATCAGGGAGGCAAGCCATGCTCA-3′ |
| | R | 5′- ACGTTGGCCCACACCACAAAGA-3′ |
| Serca2a | F | 5′-GATCCTCTACGTGGAACCTTTG-3′ |
| | R | 5′-GGTAGATGTGTTGCTAACAACG-3′ |
| Pln | F | 5′-TGCTCACTACCACATCAACTTCA-3′ |
| | R | 5′-TTCACCAAAATCAAAACTCCATTTG-3′ |

F forward, R reverse
18S was used as the household gene

**Supplemental Table I: List of primers used for reverse transcription (RT) quantitative polymer chain reaction (qPCR)**

**Supplemental Figures and Supplemental Figure legends**

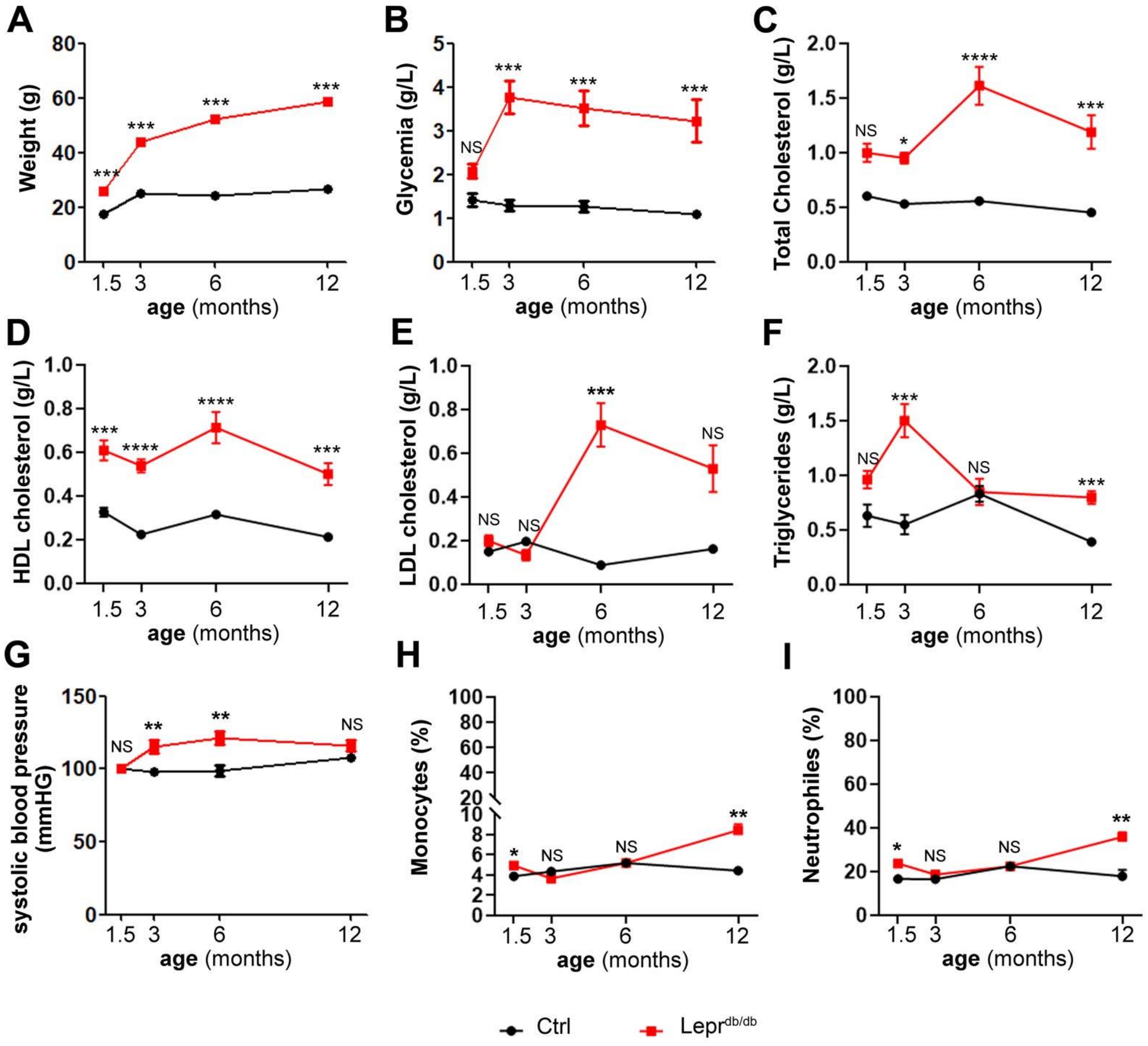


**Supplemental Figure 1: (A)** The weight of Lepr$^{db/db}$ female mice and control Lepr$^{db}$/+ mice was measured at 1, 5, 3, 6 and 12 months of age. **(B-F, H-I**) Lepr$^{db/db}$ female mice and their control Lepr$^{db}$/+ littermates were harvested with blood at 1, 5, 3, 6 and 12 months of age. Glycaemia (**B**), Total cholesterol (**C**), HDL cholesterol (**D**), LDL cholesterol (**E**) and Triglycerides (**F**) were measured. (**G**) Systolic blood pressure was measured invasively using a pressure catheter in Lepr$^{db/db}$ female mice and control Lepr$^{db}$/+ mice at 1,5, 3, 6 and 12 months of age. Circulating monocyte (**H**) and neutrophils (I) were counted. *: p≤0.05. **: p≤0.01. ***: p≤0.001. NS: not significant. Two way ANOVA followed by Bonferroni's multiple comparison test

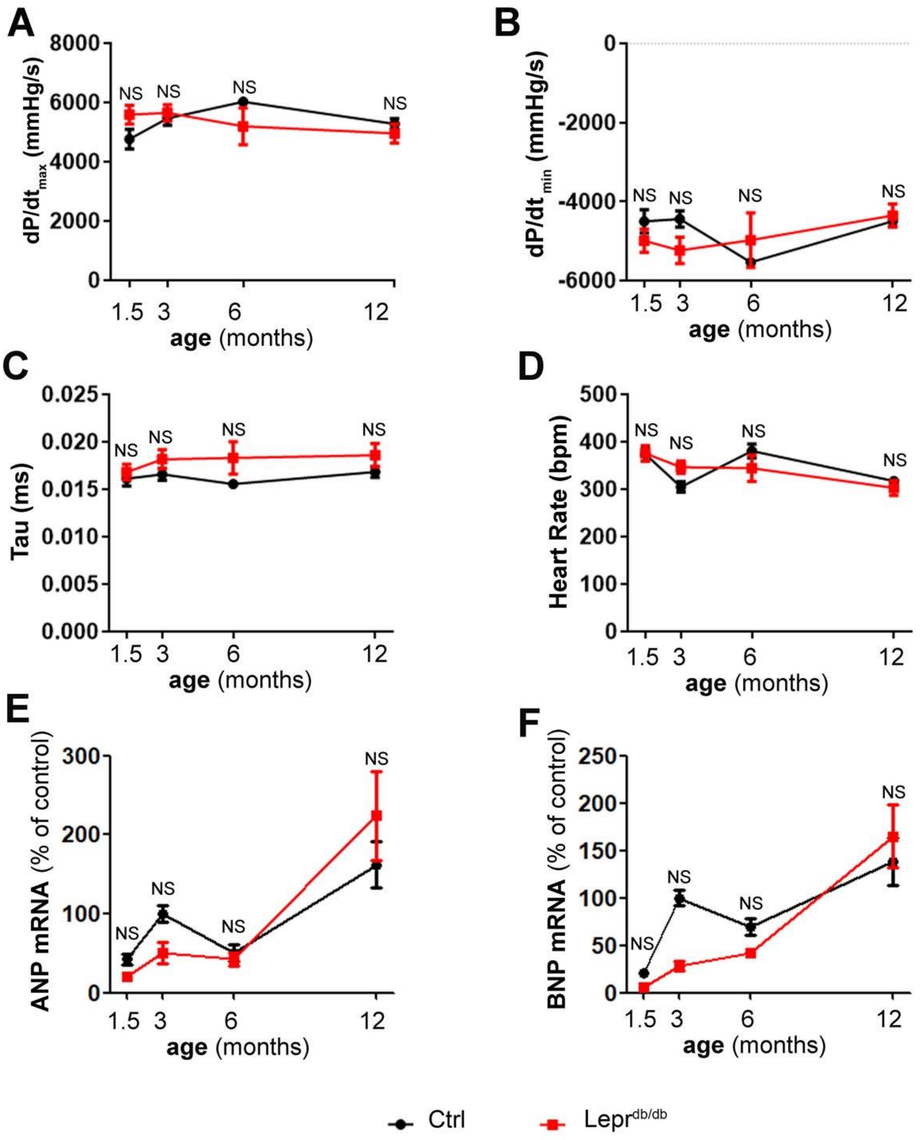


**Supplemental Figure 2:** Lepr$^{db/db}$ female mice and their control Lepr$^{db}$/+ littermates were subjected LV catheterization and sacrificed at the indicated time points (n=8 to 15 per group). dP/dt maximum (**A**), dP/dT minimum (**B**), Tau (**C**) and the heart rate (**D**) were recorded. ANP (**E**) and BNP (**F**) mRNA expression was measured via RT-qPCR in heart biopsies. NS: not significant. Two way ANOVA followed by Bonferroni's multiple comparison test.

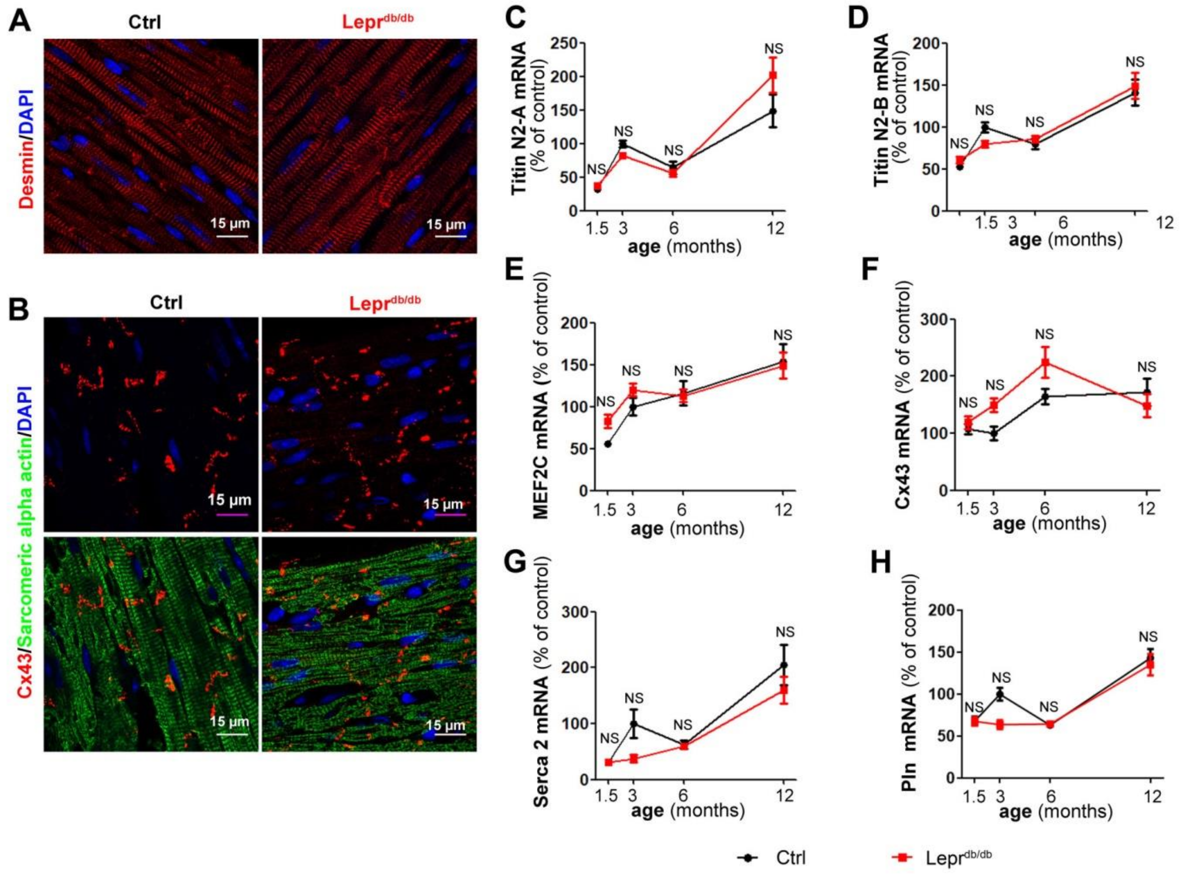


**Supplemental Figure 3:** Lepr$^{db/db}$ female mice and their control Lepr$^{db}$/+ littermates were sacrificed at 3 months of age. (**A**) Heart cross sections were immunostained with anti-Desmin antibodies to identify cardiomyocyte sarcomeres. Representative pictures are shown. (**B**) Heart cross sections were co-immunostained with anti-Cx43 antibodies (in red) to identify cardiomyocyte Intercalated discs and anti-Sarcomeric alpha actin antibodies (in green). Representative pictures are shown.. Representative pictures are shown. Titin isoform N2-A (**C**), Titin isoform N2-B (**D**), MEF2C (**E**), Cx43 (**F**), Serca2 (**G**) and Pln (**H**) mRNA expression was measured via RT-qPCR in heart biopsies. NS: not significant. Two way ANOVA followed by Bonferroni's multiple comparison test

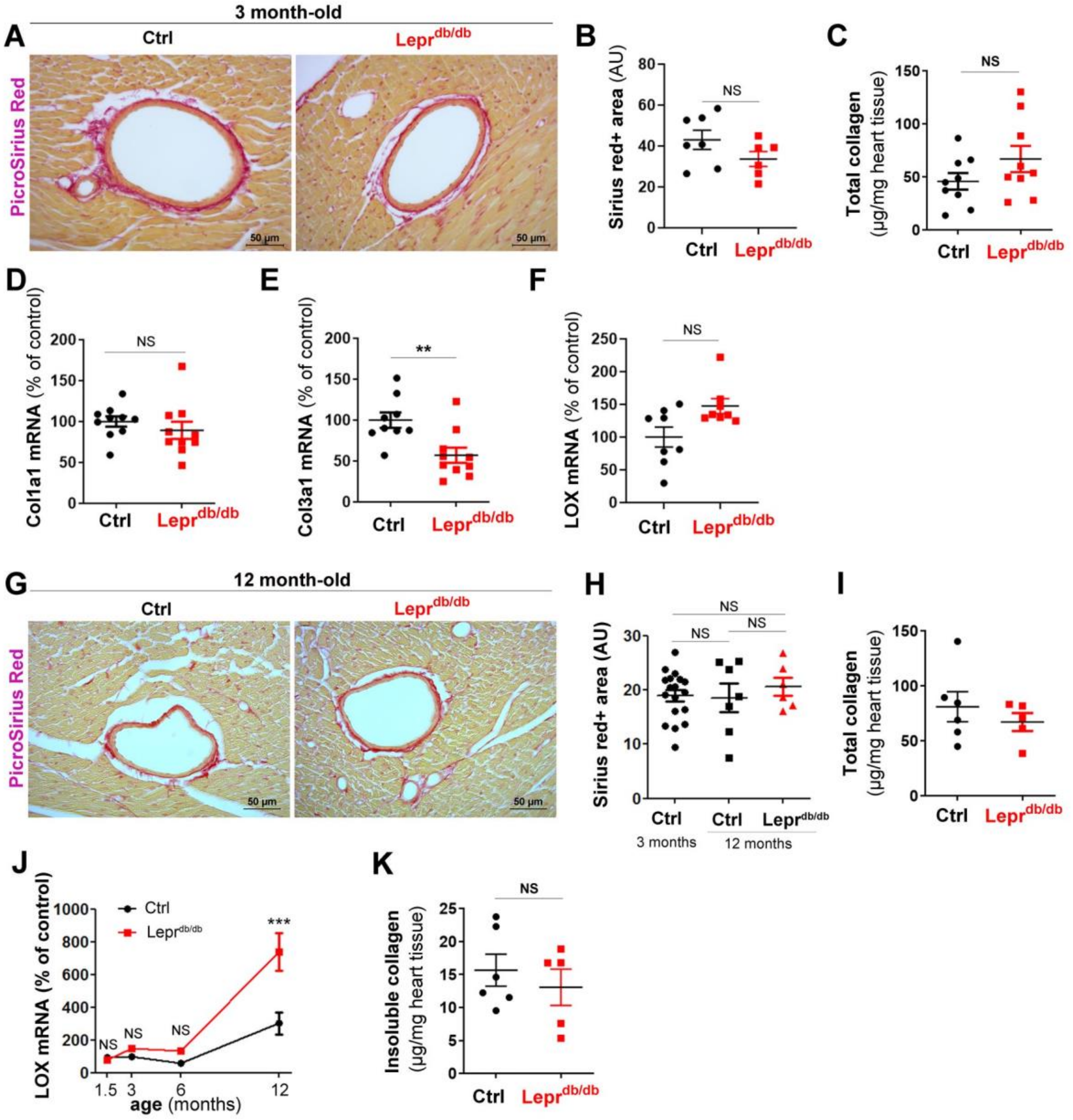


**Supplemental Figure 4: Lepr$^{db/db}$ female mice do not display significant cardiac fibrosis.** (**A-F**) Lepr$^{db/db}$ female mice and their control Lepr$^{db}$/+ littermates were sacrificed at 3 months of age. (**A**) Heart cross sections were stained with Sirius red to identify collagen. Representative pictures are shown. (**B**) Fibrosis was quantified as the Sirius red+ surface area (n=7 mice/group). (**C**) Total cardiac collagen was assessed using the Sircol™ assay. Col1a1 (**D**) Col3a1 (**E**) and LOX mRNA expression was measured via RT-qPCR in heart biopsies. (**G-I, K**) Lepr$^{db/db}$ female mice and their control Lepr$^{db}$/+ littermates were sacrificed at 12 months of age. (**G**) Heart cross sections were stained with Sirius red to identify collagen. Representative pictures are shown. (**H**) Fibrosis was quantified as the Sirius red+ surface area (n=7 mice/group). (**I**) Total cardiac collagen was assessed using the Sircol™ assay. (**J**) LOX mRNA expression was measured via RT-qPCR in heart biopsies at the indicated time points. (**K**) Unsoluble cardiac collagen was assessed using the Sircol™ assay. **: p≤0.01. ***: p≤0.001. NS: not significant. Two way ANOVA followed by Bonferroni's multiple comparison test or Mann Whitney test.

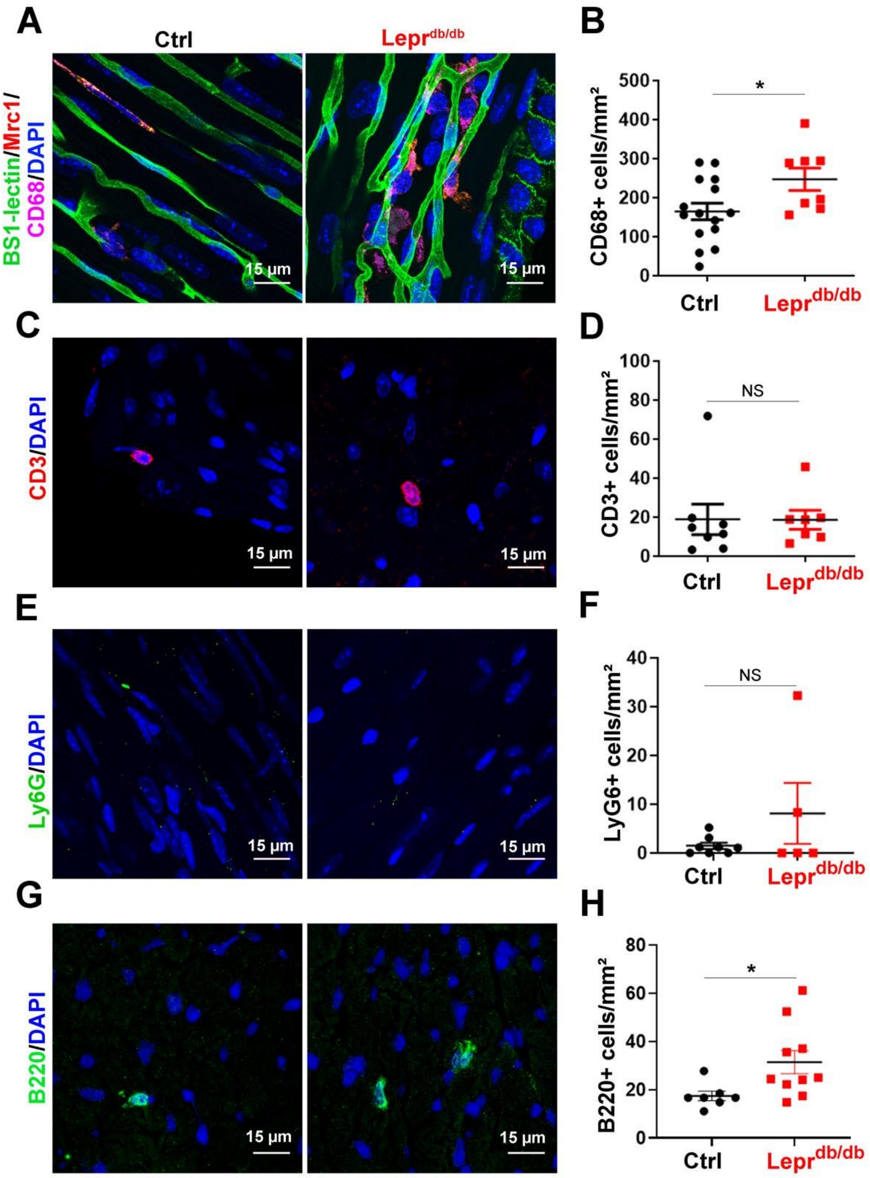


**Supplemental Figure 5:** Lepr$^{db/db}$ female mice and their control Lepr$^{db}$/+ littermates were sacrificed at 3 months of age. (**A**) Heart cross sections were immunostained with anti-CD68 antibodies to identify macrophages. Representative pictures are shown. (**B**) Macrophage infiltration was quantified as the number of CD68+ cells/mm². (**C**) Heart cross sections were immunostained with anti-Ly6G antibodies to identify macrophages. Representative pictures are shown. (**D**) Neutrophil infiltration was quantified as the number of Ly6G+ cells/mm². (**E**) Heart cross sections were immunostained with anti-CD3 antibodies to identify T Lymphocytes. Representative pictures are shown. (**F**) T-Lymphocyte infiltration was quantified as the number of CD3+ cells/mm². (**E**) Heart cross sections were immunostained with anti-CD3 antibodies to identify T Lymphocytes. Representative pictures are shown. (**F**) T-Lymphocyte infiltration was quantified as the number of CD3+ cells/mm². (**E**) Heart cross sections were immunostained with anti-B220 antibodies to identify B Lymphocytes. Representative pictures are shown. (**F**) B Lymphocyte infiltration was quantified as the number of B220+ cells/mm². *: p≤0.05. **: p≤0.01. ***: p≤0.001. NS: not significant. Mann Whitney test.

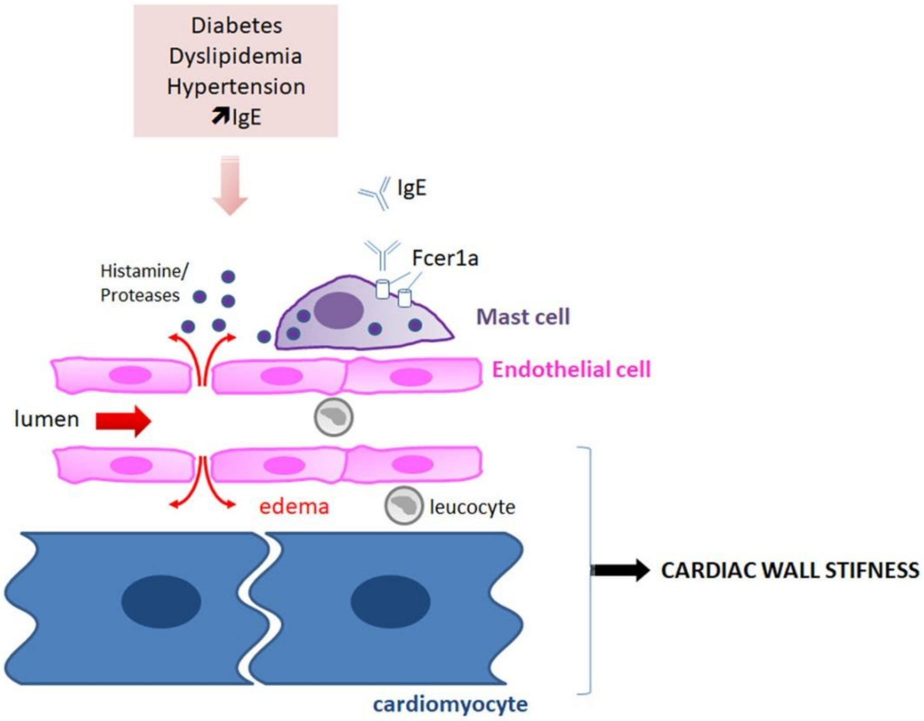


**Supplemental Figure 6:** Schema recapitulating the findings
